# Distinct contemplative traditions share a common EEG complexity signature

**DOI:** 10.1101/2025.06.20.660652

**Authors:** Prakash Shrimali, Arun Sasidharan, Saketh Malipeddi, Bianca Ventura, Rahul Venugopal, Ajay Kumar Nair, Prejaas K.B. Tewarie, P.N. Ravindra, Bindu M Kutty, Georg Northoff

**Affiliations:** Centre for Consciousness Studies (CCS), Department of Neurophysiology, Basic Science Block, National Institute of Mental Health and Neuro Sciences (NIMHANS), Bengaluru, 560029, Karnataka, India; The Royal’s Institute of Mental Health Research & University of Ottawa, Brain and Mind Research Institute, Centre for Neural Dynamics, Faculty of Medicine, University of Ottawa, 145 Carling Avenue, Rm. 6435, Ottawa, ON K1Z 7K4, Canada; Department of Cellular and Molecular Medicine, University of Ottawa, Ottawa, ON, Canada; Centre for Brain and Mind (CBM), Department of Psychiatry, National Institute of Mental Health and Neuro Sciences (NIMHANS), Bengaluru, Karnataka, India; Joint International Research Unit on Consciousness, CERVO Brain Research Centre, University of Laval, Québec City, G1J 2G3, Canada; Sir Peter Mansfield Imaging Centre, School of Physics, University of Nottingham, Nottingham, NG7 2RD, United Kingdom

## Abstract

Meditation encompasses diverse practices that train attention inward, unlike externally oriented task states. Which brain-activity features are shared across contemplative practices, which accompany long-term training, and which are tradition-specific remains unclear. We leveraged electroencephalography data from 226 participants: advanced meditators of four contemplative traditions and non-meditating controls. We developed a neurodynamic profiling approach spanning 21 features across four feature families: oscillatory, aperiodic, complexity and timescale. Using extensive machine learning, we found that meditation and task states were reliably separable, driven by a large, uniform drop in complexity during meditation. Meditators differed from controls primarily in timescale dynamics. Traditions exhibited distinct signatures in all feature families except complexity, and meditators’ neurodynamic profiles identified the contemplative tradition they practiced. A classifier trained on three traditions could still detect meditation in the fourth, regardless of which tradition was left out, relying mainly on complexity features. Diverse contemplative techniques therefore converge on a shared neurodynamic signature—reduced complexity during meditation relative to an external task, that generalizes across traditions. This shared signature coexists with tradition-specific profiles in oscillatory, aperiodic, and timescale features.

## Introduction

For millennia, meditation traditions have engineered diverse practices that differ in technique. For instance, in the Brahma Kumaris tradition, practitioners visualise a point of light between the brows. In Heartfulness, they absorb on a felt centre at the heart. In Isha, they observe the movement of the breath, while in Vipassana, attention is swept through the body. That is, meditation comprises a diverse range of practices for cultivating attention and awareness, typically in the service of self-regulation ^1–3^. The traditions converge on a similar goal: liberation from suffering by loosening automatic identification of the self with the contents of experience, i.e. de-reification ^1,4^. We therefore raise the question whether such phenomenological similarity across different traditions relates to an analogous neurodynamic similarity in the brain mediating such internal attentional state shared across diverse meditative techniques. Do diverse internally oriented techniques, despite differences in attentional focus and phenomenology, converge on a common neurodynamic signature when contrasted with externally oriented attention?

What these diverse techniques share is a common orientation in their attention: attention is directed inward, toward the interoceptive or the self-related psychological aspects, characterizing meditation as an internally oriented state ^5–7^. Meditation being internally oriented contrasts with the externally oriented attentional focus typical of non-meditative states like task-based states, in which attention is directed outward toward visual, auditory, or other environmental stimuli ^8,9^. Internally and externally oriented states reflect distinct forms of cognition ^10,11^, although they may arise as different internal–external balances from a shared underlying baseline ^9^. Whether these diverse internally oriented meditation practices share common neural signature that distinguish them from externally oriented task state remains an open question.

Based on this, we asked which neurodynamic features carry the signature that distinguishes the internally oriented meditative from externally oriented task state. Prior work has examined topographic differences between default-mode and central-executive networks during meditation ^12^, but the temporal and thus the neurodynamic signatures of internal versus external attentional states remain poorly understood, particularly across different meditation traditions. Electroencephalography studies have reported a general slowing of rhythms during meditation ^13^, increases in alpha and theta power linked to inward attention ^14^, and increases in gamma linked to perceptual integration ^15,16^. These oscillatory findings are inconsistent across studies, however, and some appear technique-specific ^17^, with limited evidence that they represent markers of a meditative state shared across practices. Oscillatory changes have also been reported during intracranial recordings of loving-kindness meditation, without corresponding changes in aperiodic activity ^18^. A characterization of the shared neural dynamics across meditation traditions may require looking beyond the widely studied oscillatory measures, though ^14,17^.

Three lines of evidence support a broader neurodynamic approach reaching beyond oscillations. First, meditation is associated with changes in signal complexity, particularly in experienced practitioners ^19–21^. Importantly, the direction of these effects depends on the comparison being made: complexity may increase during meditation relative to waking rest or mind-wandering, while experience practitioners may also exhibit lower baseline complexity as a trait of practice ^19,22^. This distinction is particularly relevant given our use of an externally oriented task as the comparison condition, instead of rest. Second, parameterizing the power spectrum into oscillatory and aperiodic components reveals meditation-related changes in the aperiodic 1/f slope which has been linked to excitation–inhibition balance ^23,24^. The slope has been reported to steepen during meditation in experienced but not novice practitioners ^20^.

Third, intrinsic neural timescales, indexed by the autocorrelation window, are modulated by meditation experience, attentional orientation, and meditation technique ^25–28^. Together, these findings suggest that meditation-related neural dynamics extend across multiple dynamic properties of the neural signal that are not fully captured by oscillatory measures alone. We therefore adopted a neurodynamic profiling approach by assembling 21 features spanning four categories — oscillatory activity, aperiodic activity, signal complexity and intrinsic neural timescales (Table 1 in methods). These features were analysed jointly using machine learning approach to identify neurodynamic patterns that are shared across, or differentiate between, meditation traditions.

**Table 1.**
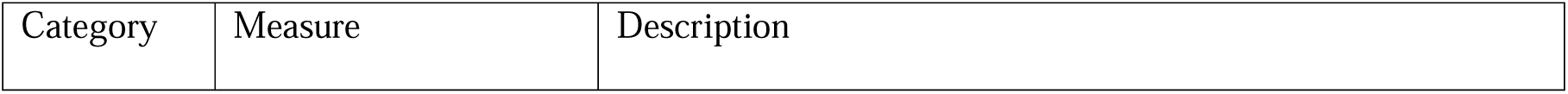
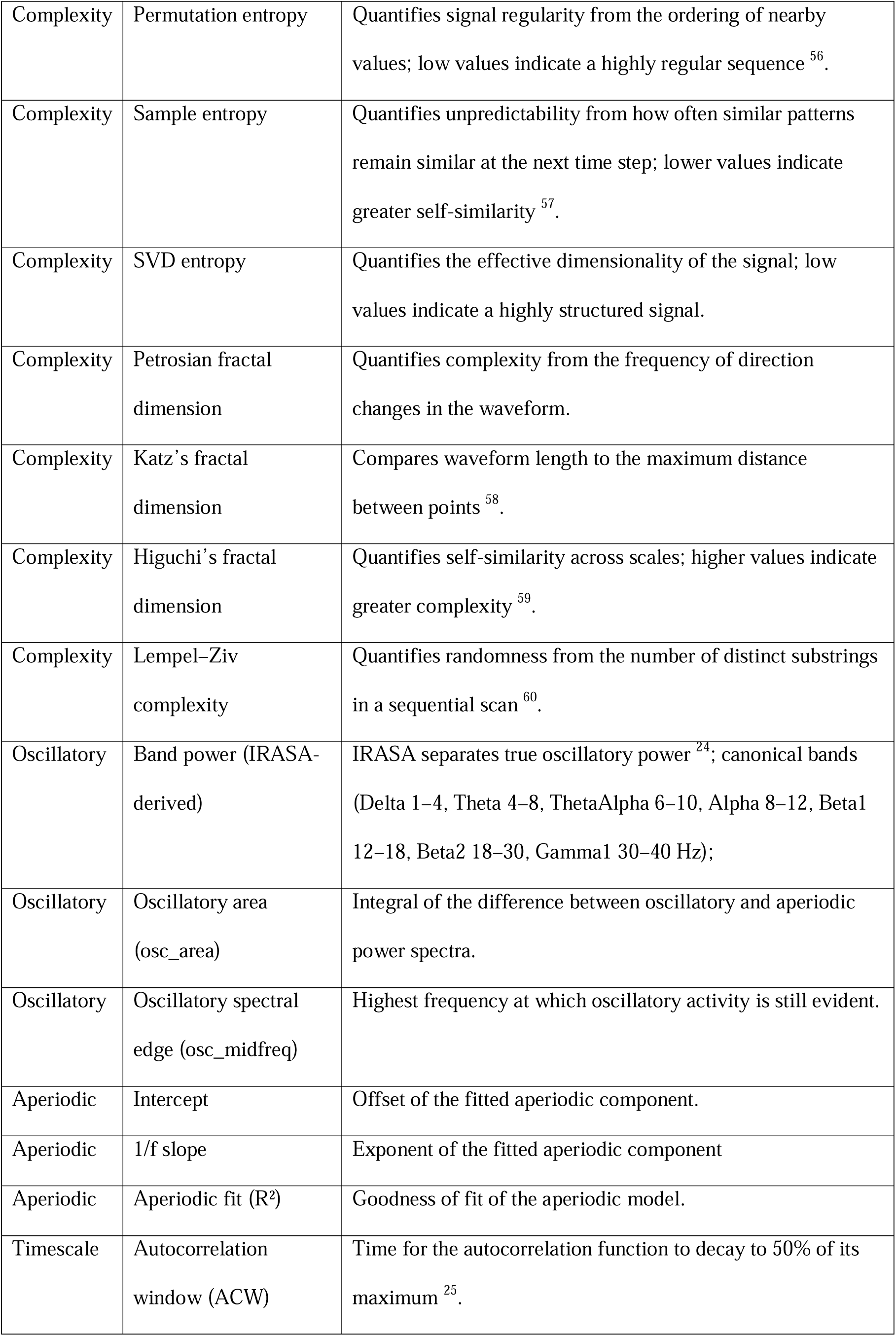
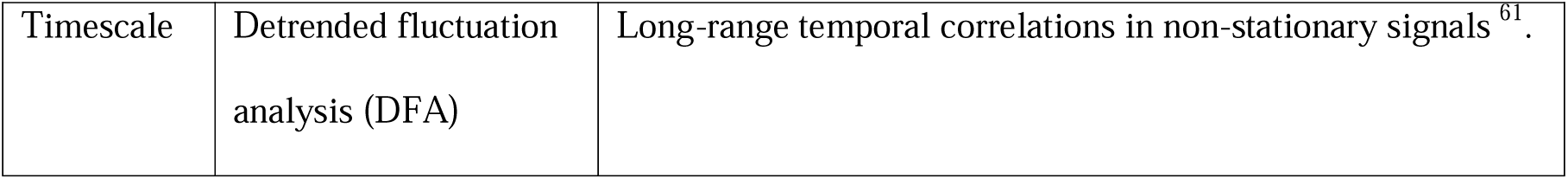
Neurodynamic measures included in the present study.

Progress, however, has been constrained not only by limited neurodynamic feature repertoire but also the structure of the available data. The ENIGMA-Meditation consortium recently characterized the field as relying largely on small datasets that are heterogeneous in practice type, experience level, study design and imaging modality, with population-level generalizability yet to be established ^29^.

Meta-analytic evidence points to related limitations, including: moderate effect sizes and concerns about publication bias ^30^. Moreover, most studies examine a single practice or aggregate practices, limiting the ability to distinguish neural features that are shared across traditions from those that are practice-specific ^31^. What lacks is a larger-scale dataset in which multiple meditation traditions are assessed against a common comparator (such as an external task), using the same recording system and preprocessing pipeline. Such a dataset is necessary to distinguish those neurodynamic features that are shared across practices from those that are tradition-specific, a distinction that cannot be made by aggregating heterogeneous studies or examining single traditions in isolation. Our study addresses this gap by combining such large-scale dataset with a multidimensional neurodynamic profile to investigate which neurodynamic features are shared across meditation traditions and which differentiate them.

Here we analysed EEG from 226 participants, comprising long-term practitioners of Vipassana, Isha Yoga, Heartfulness and Brahma Kumaris Raja Yoga and non-meditating controls, recorded during internal meditation and a common external task condition. We used extensive machine learning jointly with 21 features (see Table 1) to address three questions, ordered from the most general to the most specific:

1. *State*: which neurodynamic features distinguish meditation from task across techniques and groups, and do these features generalize across traditions?
2. *Experience*: does long-term practice change which neurodynamic features carry the distinction?
3. *Tradition*: Which neurodynamic features differentiate the four traditions from one another, and which generalize across them?

We define a ’shared meditative state’ operationally as a neurodynamic pattern that (1) distinguishes meditation from the external task and (2) generalizes to a tradition that was not represented during classifier training. This definition is stricter than within-tradition accuracy, as high within-tradition accuracy could arise from four separate, technique-specific solutions.

## Results

Throughout the text, Med denotes the meditation condition and Task the externally oriented task condition.

### A reproducible multivariate meditation–task distinction

We first asked whether meditation can be told apart from the external task on the basis of the EEG features alone, without any knowledge of the practice a participant follows. Pooling all participants and using all measures from Table 1 (complexity, oscillatory aperiodic and timescale), a Random Forest classifier separated Med from Task with a mean balanced accuracy of 0.809 (SD 0.036) and mean area under curve (AUC) of 0.887 (SD 0.024) across 100 repeated subject-held-out splits (Fig. 2a,b).

**Figure 1.**
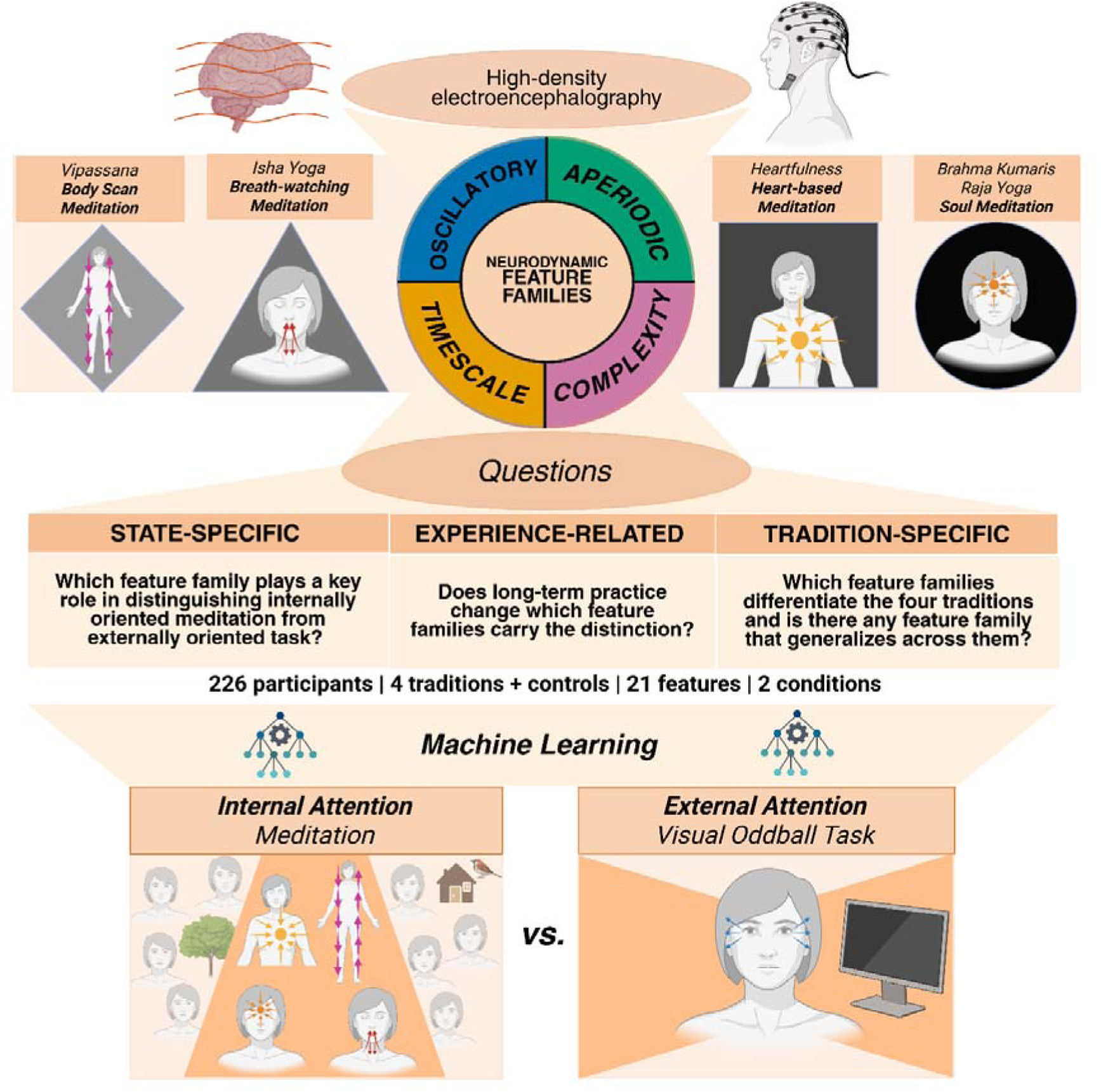
Study design and analytic framework. EEG was recorded with 128-channel nets from 226 participants: long-term practitioners of four contemplative traditions (Vipassana body-scan, Isha Yoga breath-watching, Heartfulness heart-based, Brahma Kumaris Raja Yoga soul meditation) and non-meditating controls — during meditation and during a visual oddball task requiring externally directed attention. Each recording was summarised by 21 measures spanning four neurodynamic feature families: oscillatory, aperiodic, complexity and timescale. Machine-learning analyses then addressed three questions: which families distinguish internally oriented meditation from the externally oriented task; whether long-term practice changes which families carry that distinction; and which families differentiate the four traditions, and whether any generalises across them.

**Fig 2.**
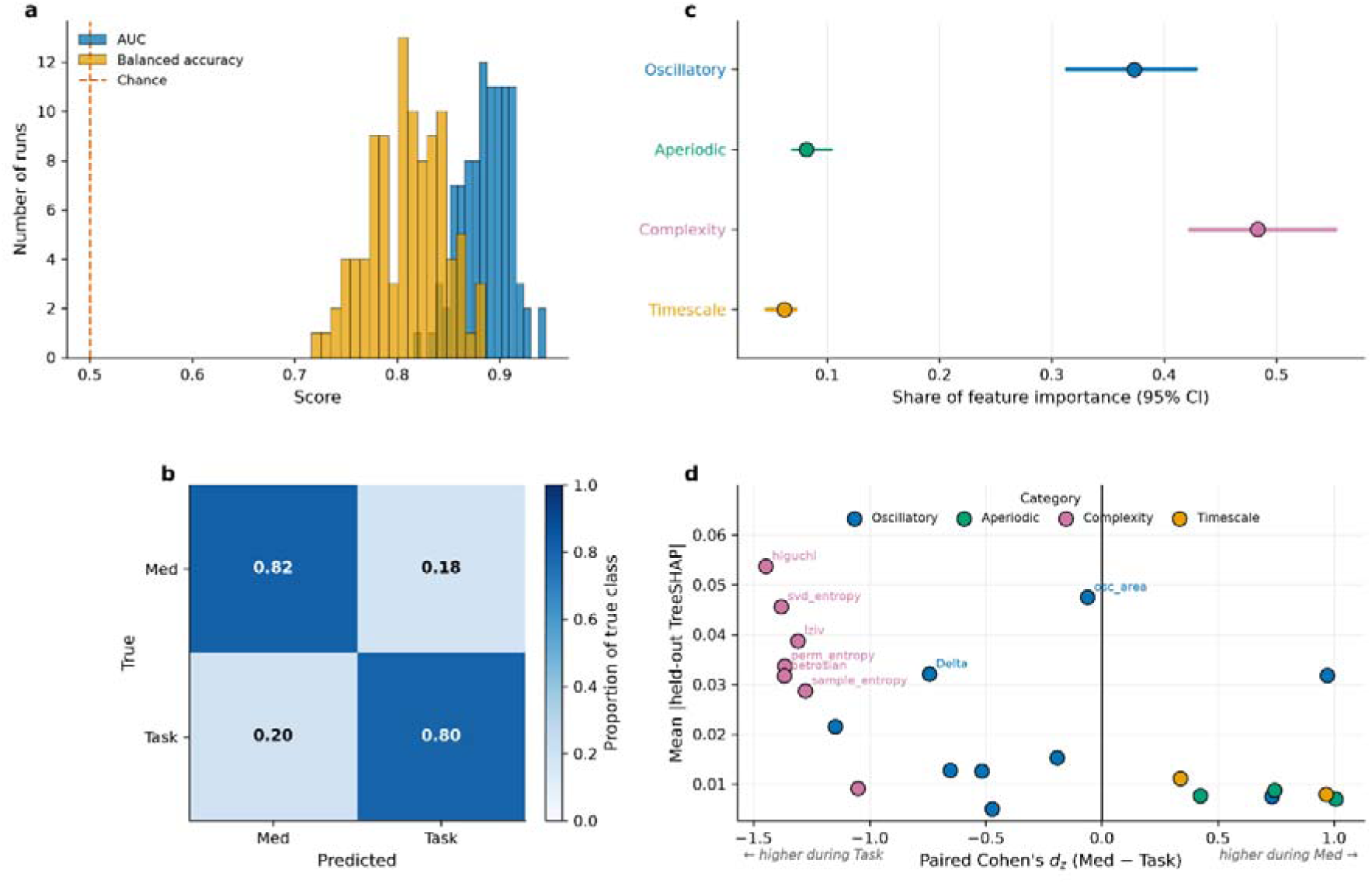
A reproducible multivariate distinction between meditation and task states. **a,** Distribution of AUC and balanced accuracy across 100 repeated subject-held-out splits; dashed line marks chance. **b,** Mean confusion matrix, normalized within each true class. **c,** Share of Random Forest feature importance per neurodynamic category; points are estimates and lines 95% confidence intervals from bootstrap resampling of subjects. **d,** Each point is one feature, positioned by its paired Med−Task effect size (x) and mean absolute held-out TreeSHAP value (y). Filled points indicate FDR-adjusted *P*<0.05 for the Med−Task change. Position on x gives the direction and size of the physiological change; height on y gives the contribution to classification.

Feature-level effects were large and highly consistent. All 21 features showed a significant paired Med−Task change (FDR *q*<0.05), and the largest effects were complexity measures, every one of which was lower during meditation than during the task: Higuchi’s fractal dimension (*d*z=−1.45), SVD entropy (−1.38), Petrosian fractal dimension (−1.37), permutation entropy (−1.37), Lempel–Ziv complexity (−1.31), sample entropy (−1.28) and Katz’s fractal dimension (−1.05), with a mean effect of *d*z=−1.31 (negative values denote lower values during meditation than during the task). The aperiodic slope steepened (+1.01), and alpha power (+0.97) and both timescale measures (mean +0.65) increased, while Beta2 power decreased (−1.15). Taken together, the EEG during meditation was more regular and more predictable than during the task, with power shifted from fast beta activity towards alpha and a steeper fall-off of power with frequency. Longer timescales indicate that the signal remained correlated with its own past for longer, while lower complexity in this sense describes a less irregular signal in meditation.

Complexity also carried the largest share of classifier importance (0.483, 95% CI [0.423, 0.553]), ahead of oscillatory (0.373 [0.313, 0.429]), aperiodic (0.082 [0.070, 0.103]) and timescale (0.062 [0.046, 0.072]) features (Fig. 2c). The confidence intervals for complexity and oscillatory features overlap, so this difference should be interpreted cautiously. The redundancy-adjusted shares, which account for intercorrelation among features, show complexity leading more clearly (0.668, Supplementary Fig. S1). Complexity contains more measures than the other categories, and those measures are strongly intercorrelated (mean |ρ|=0.89; 1.21 effective dimensions from seven measures). We therefore repeated the comparison under two normalizations. Complexity retained the largest share under raw (0.500), per-feature (0.429) and redundancy-adjusted (0.668) weighting (Supplementary Fig. S1), so its leading contribution is not explained by the number or redundancy of its measures. Within each group and each tradition separately, complexity also ranked first once redundancy was considered, although in the control, Isha Yoga and Vipassana cohorts the raw shares placed oscillatory features first (Supplementary Fig. S1b).

Plotting predictive importance against physiological change separates two distinct quantities (Fig. 2d): complexity measures occupy the region of high classifier importance and strongly negative Med−Task change, whereas the aperiodic slope and alpha power contribute with the opposite sign.

Together, these analyses establish the first result: meditation can be reliably distinguished from the externally oriented task, with complexity showing the largest contribution and a consistent decrease during meditation. The next question is whether this distinction depends on long-term meditation experience.

### The Med-Task distinction is not identical in practitioners and controls

Meditators and controls showed reliable classification (advanced practitioners 0.847, AUC 0.913; controls 0.803, AUC 0.888), and the direction of the complexity effect was the same in both (mean *d*z=−1.40 and −1.19 respectively). In both groups, complexity was lower during meditation than during the task. The state distinction is therefore not a property of expertise as it is also present in people meditating for the first time.

At the category level, the two groups differed reliably in only one family: timescale features contributed less in practitioners than in controls (0.040 versus 0.080, difference −0.039; permutation *P*=0.003, *q*=0.012). In other words, the classifier relied only about half as much on timescale features to recognize meditation in practitioners as it did in controls. The differences in complexity (0.527 versus 0.400, *P*=0.066) and oscillatory contributions (0.330 versus 0.423, *P*=0.147) were in the expected directions but not statistically supported at this sample size, and the aperiodic contribution was almost identical in the two groups (0.103 versus 0.098, *P*=0.589; Fig. 3a). Repeating these tests on importance expressed per effective dimension again identified only the timescale difference (0.053 versus 0.113, *P*=0.002), with complexity closer to, but still short of, the threshold (0.687 versus 0.565, *P*=0.039, *q*=0.077; Supplementary Fig. S1c).

**Fig 3.**
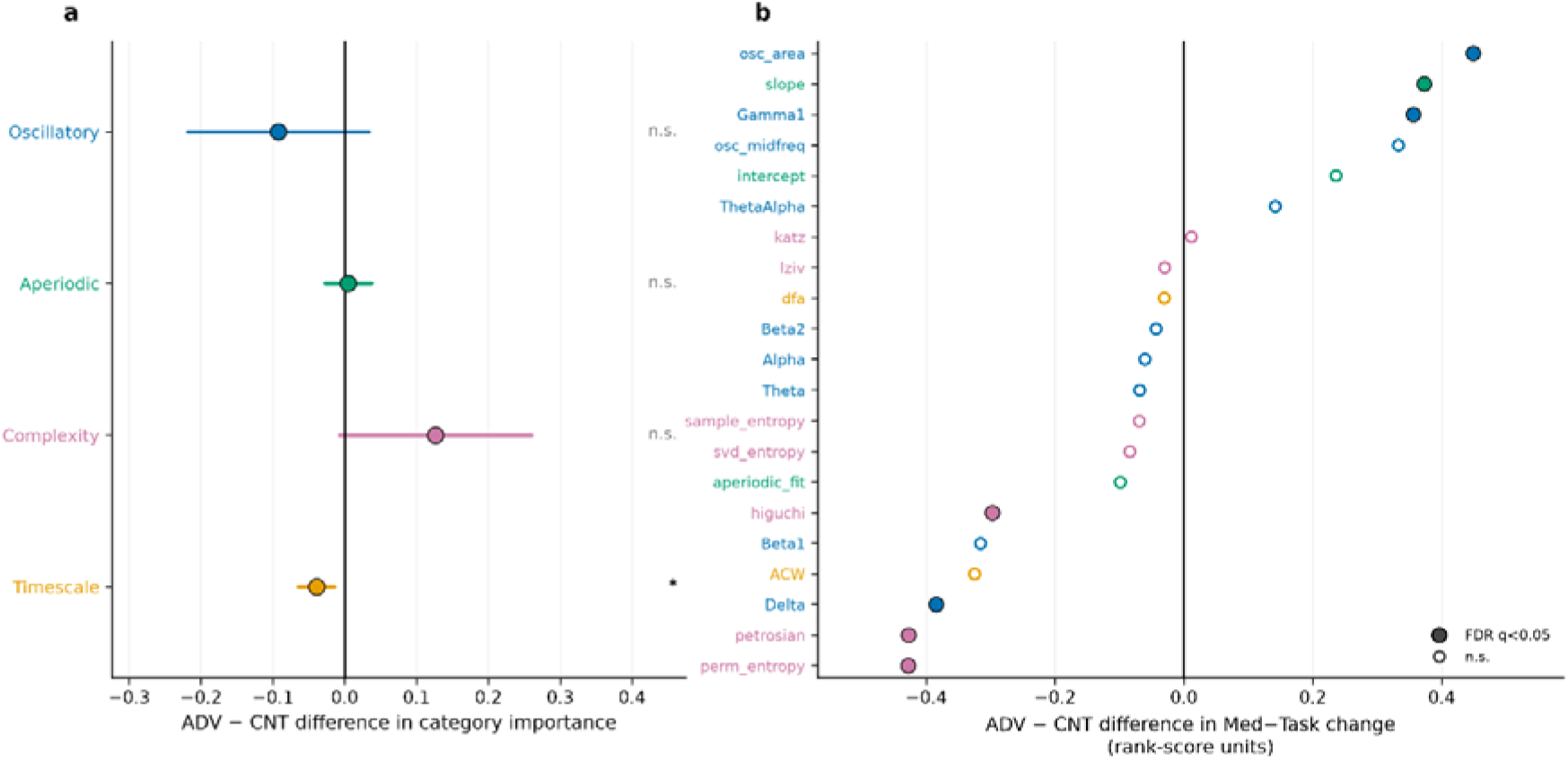
The meditation–task profile differs between advanced practitioners and controls. **a,** Difference between practitioners and controls in category importance; points are observed differences and lines span 1.96 SD of the permutation null. **b,** Feature-level Group × Condition effects from age-adjusted mixed-effects models, as the between-group difference in estimated Med−Task change. Filled points indicate FDR-adjusted *P*<0.05.

Individual features were more sensitive to group differences than whole categories. For each feature, an age-adjusted mixed-effects model tested whether the Med−Task change differed between practitioners and controls, which is a Group × Condition interaction. This was the case for 7 of 21 features (FDR *q*<0.05): three complexity measures (permutation entropy, LR(1)=12.6, *q*=0.004; Petrosian fractal dimension, LR(1)=12.6, *q*=0.004; Higuchi’s fractal dimension, *q*=0.033), three oscillatory measures (Gamma1, oscillatory area and Delta) and the aperiodic slope (*q*=0.018; Fig. 3b). For the complexity measures, the decrease during meditation was larger in practitioners than in controls. Long-term practice thus deepens the shift towards a more regular signal that controls already show, without changing its direction. The state distinction is therefore present in both groups, while long-term practice changes the magnitude and relative contribution of some features. We next asked whether the meditation–task profile can also distinguish the four traditions from one another.

### Tradition identity is recoverable from an individual’s profile

We next compared the four traditions with one another and excluded controls throughout, since the question here concerns differences between techniques and the practitioner–control contrast was addressed above. If the four practices produce genuinely different neurodynamic profiles, it should be possible to recover which practice a participant follows from their own Med−Task change alone, i.e., from the difference between each feature during meditation and during the task. A four-class classifier decoded tradition identity with 0.620 balanced accuracy (95% CI [0.613, 0.627]) against a permutation-derived chance level of 0.245 (*P*<0.001; Fig. 4a), so roughly six in ten practitioners were assigned to their own tradition, compared with one in four expected by chance. Adding age improved this modestly to 0.657. Tradition identity is therefore recoverable, although far less reliably than the meditation–task distinction itself (0.620 versus 0.809). Each technique leaves a measurable imprint on the signal, but that imprint is weaker than the shift shared by all practice-the two-tiers.

**Fig 4.**
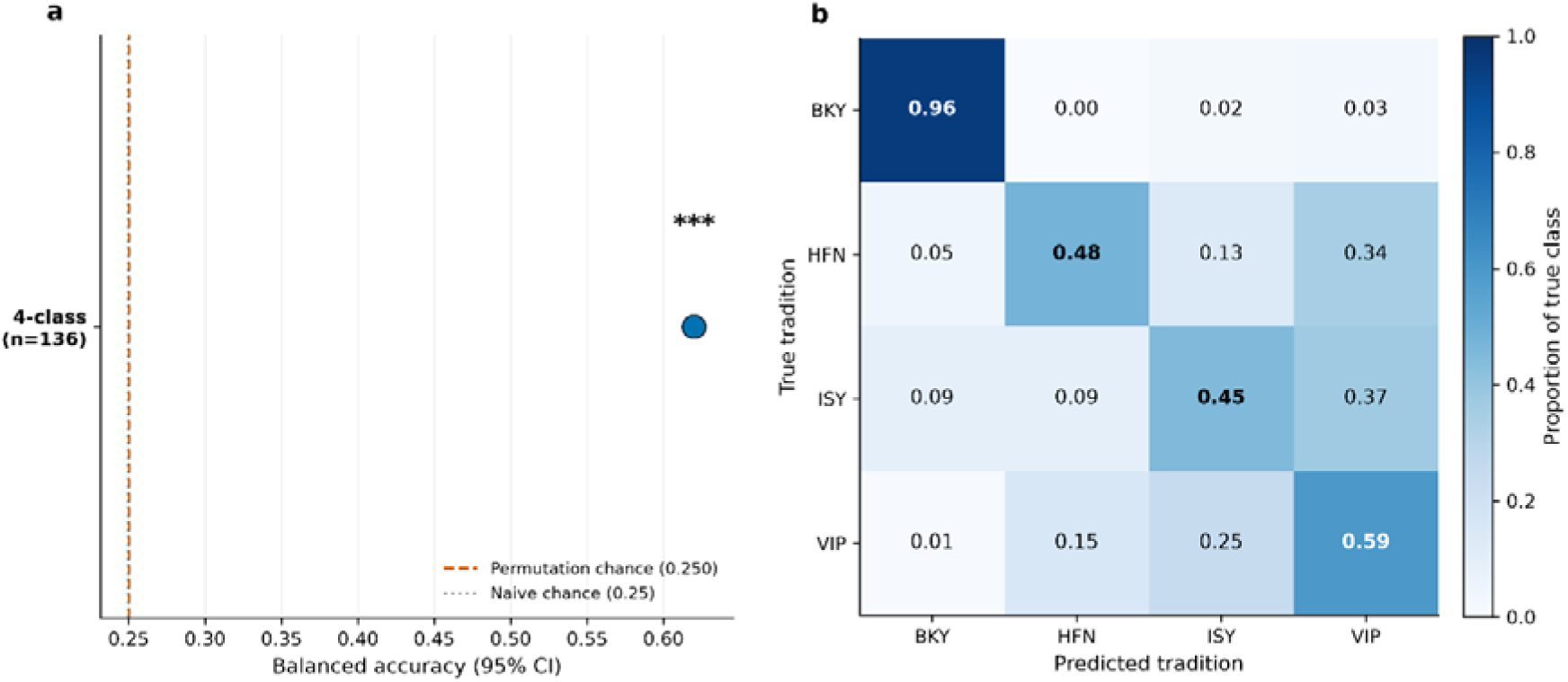
Tradition identity is recoverable from an individual’s meditation–task profile. **a,** Balanced accuracy of a four-class ensemble classifier decoding tradition from subject-level Med−Task profiles. Dashed line is the permutation-derived chance level; dotted line the naive 1/k value. **b,** Out-of-fold confusion matrix, normalized within each true tradition.

The confusion matrix shows the accuracy to be unevenly distributed, and its errors to be informative (Fig. 4b). Brahma Kumaris Raja Yoga was recovered almost perfectly (recall 0.957), whereas the other three were recovered at 0.446–0.595, with the principal confusions between Isha Yoga and Vipassana (0.370 and 0.245 in the two directions) and from Heartfulness to Vipassana (0.345). The two breath- and body-focused practices are thus the pair most easily mistaken for one another.

The ability to recover tradition identity shows that the four practices leave distinguishable imprints on the meditation–task profile. We next asked which feature families account for these differences, and whether any component remains common across all four traditions.

### Traditions differ in feature families, but share a decrease in complexity

Meditation was decoded from task within every tradition (balanced accuracy: Brahma Kumaris Raja Yoga 0.968; Vipassana 0.823; Isha Yoga 0.782; Heartfulness 0.741).

We then asked whether the classifier relied on the same feature families in every tradition, by testing whether the importance of each family varied across traditions more than expected under a permutation null. Three of the four families differed in their contribution: oscillatory (spread 0.372 versus a null of 0.151, *P*=0.0018), aperiodic (0.178 versus 0.057, *P*=0.0014) and timescale (0.072 versus 0.026, *P*=0.005), all at *q*=0.0028. The contribution of complexity, in contrast, did not vary reliably across traditions (0.250 versus 0.157, *P*=0.098; Fig. 5a). This null is not an artefact of complexity being the largest family, with seven of the 21 measures: with importance expressed per effective dimension the same three families differed (oscillatory *P*=0.005, aperiodic *P*=0.005, timescale *P*=0.006) and complexity still did not (0.152 versus a null of 0.128, *P*=0.302; Supplementary Fig. S1c). Hence, each tradition weighted oscillatory, aperiodic and timescale features differently, whereas the weight given to complexity stayed relatively comparable across all four.The traditions separated the families that differed in ways that tracked the structure of each practice. Oscillatory contributions were highest in the two breath- and body-directed practices, Isha Yoga (0.518) and Vipassana (0.505), intermediate in Heartfulness (0.346), and lowest in Brahma Kumaris Raja Yoga (0.146). Aperiodic contributions ran in the opposite direction, with the highest in Brahma Kumaris Raja Yoga (0.229) and the lowest in Vipassana (0.052). Timescale contributions, small everywhere, were largest in Heartfulness (0.082) and smallest in Brahma Kumaris Raja Yoga (0.010). Complexity, by contrast, was the leading or second contributor in every tradition (0.365–0.615) without varying reliably between them. Once expressed per effective dimension, it led in all four (0.562–0.627).

**Fig 5.**
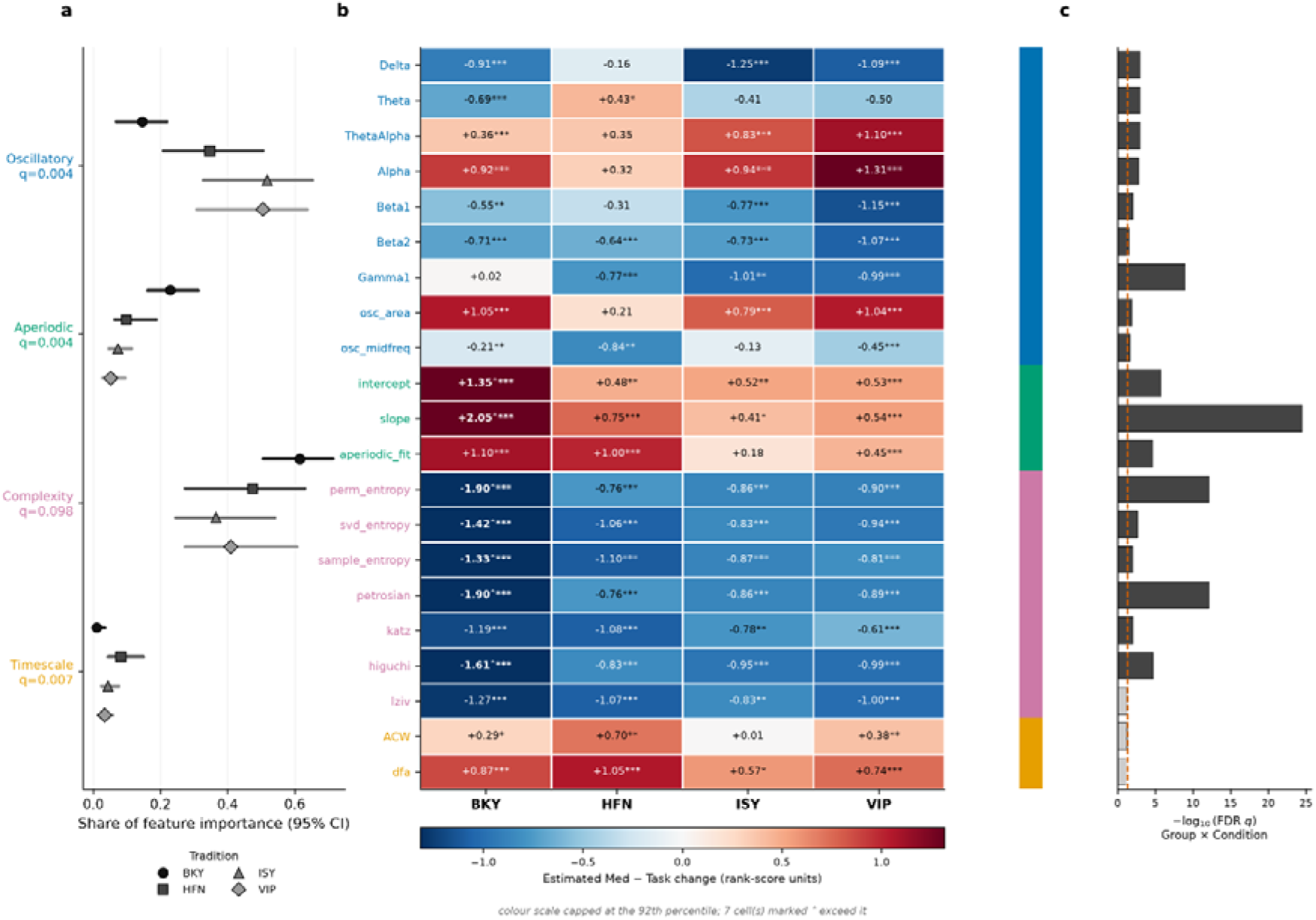
Traditions differ in feature-family contributions and in their meditation–task profiles. **a,** Category importance per tradition; points are estimates and lines 95% bootstrap confidence intervals. The *q* value beside each category is from a permutation test of whether that category’s contribution varies across traditions. **b,** Age-adjusted estimated Med−Task change per feature per tradition. Asterisks denote the within-tradition Med−Task effect, not a difference between traditions. **c,** FDR-adjusted significance of the Group × Condition interaction, which does test differences across traditions (18 of 21 features). * q<0.05, ** q<0.01, *** q<0.001.

At the level of individual features, the four traditions differed almost everywhere. An age-adjusted Group × Condition model restricted to the four traditions showed that the Med−Task change differed across traditions for 18 of 21 features (FDR *q*<0.05), most strongly for the aperiodic slope (LR(3)=123.3, *q*=3.2×10□²□), followed by Petrosian fractal dimension and permutation entropy (both LR(3)=63.5, *q*=7.6×10□¹³), Gamma1 (LR(3)=48.1) and the aperiodic intercept (LR(3)=32.6). Only three features changed similarly in all four traditions: Lempel–Ziv complexity, the autocorrelation window and detrended fluctuation analysis, all of which index the temporal organization of the signal (Fig. 5b,c).Underneath this heterogeneity, the direction of the complexity effect was invariant. All seven complexity measures decreased during meditation in all four traditions, with mean effects of *d*z=−2.71 (Brahma Kumaris Raja Yoga), −1.45 (Heartfulness), −1.34 (Vipassana) and −0.91 (Isha Yoga): the magnitude varied up to threefold, but the sign never changed. For the other families, by contrast, the size of the change depended strongly on the practice. The aperiodic slope steepened most in Brahma Kumaris Raja Yoga (estimated Med−Task +2.05) and least in Isha Yoga (+0.35). Both timescale measures lengthened most in Heartfulness (mean *d*z=+1.01) and were essentially unchanged in Isha Yoga (+0.22, with only one of the two reaching significance). Alpha power showed its largest increase in Vipassana (estimated Med−Task +1.31), where the adjacent theta–alpha band also contributed strongly. The traditions therefore differ substantially in the contribution and magnitude of several neurodynamic features, but the direction of the complexity change is shared across all four. This suggests a candidate shared component, but it does not yet establish that the shared component is sufficient to generalize across traditions.

### The distinction generalizes to an unseen tradition

The four techniques differ in most features, so a shared state can only be claimed if it survives transfer from one tradition to another. We therefore trained classifiers on three traditions and tested them on the fourth, which the model had never seen. This is a stronger test than accuracy within each tradition, because four high within-tradition accuracies could also arise from four entirely separate, technique-specific solutions. All four traditions generalized above chance (Fig. 6a): Brahma Kumaris Raja Yoga 0.886 (95% CI [0.826, 0.946], *P*<0.001), Vipassana 0.852 ([0.783, 0.919], *P*<0.001), Isha Yoga 0.821 ([0.742, 0.903], *P*<0.001) and Heartfulness 0.707 ([0.591, 0.818], *P*<0.001), against permutation nulls centred near 0.50. A model that had never encountered a practice therefore still classified at least seven in ten recordings from that practice correctly. Confidence intervals are from bootstrap resampling of held-out subjects, and *P* values from permuting condition labels within subject in the training set.

**Fig 6.**
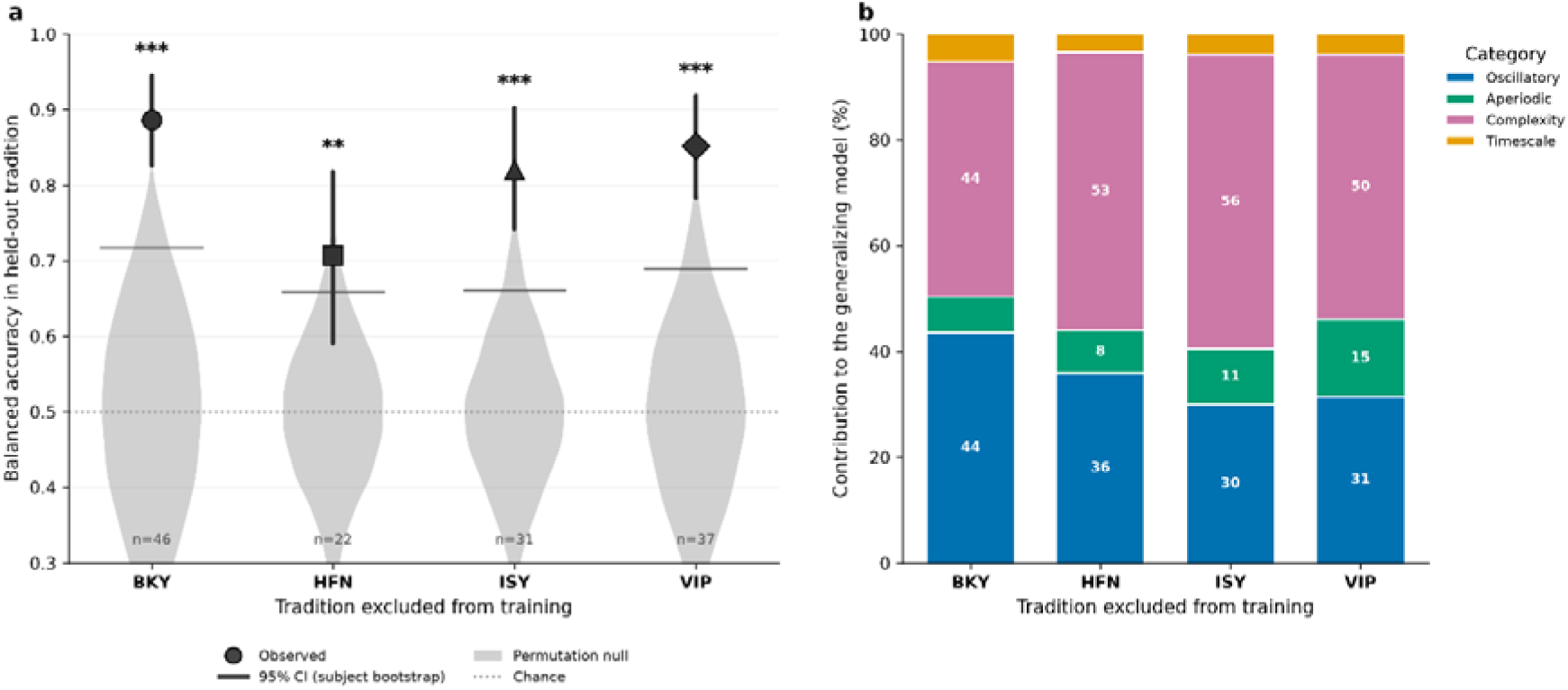
Cross-tradition generalization of the meditation–task distinction. **a,** A classifier trained on three traditions and tested on the fully held-out fourth. Points are balanced accuracy, lines 95% confidence intervals from bootstrap resampling of held-out subjects, and shaded distributions the null from permuting condition labels within subject in the training set. Dotted line marks chance. \**P*<0.05, \*\**P*<0.01, \*\*\**P*<0.001. **b,** Composition of the generalizing models as the percentage of feature importance per category. Shares are unadjusted, matching Figs. 2c, 3a and 5a; the redundancy-adjusted version is in Supplementary Fig. S1b.

Complexity was the leading contributor in every held-out model, carrying 0.443–0.557 of feature importance (Fig. 6b). This holds even where complexity was not the leading contributor within the held-out tradition itself: for Isha Yoga, whose own profile is oscillatory-dominated (0.518), the model that successfully classified it was complexity-dominated (0.557). The signature that transfers between traditions is therefore carried by complexity, even for a practice whose own profile is dominated by oscillatory features. The distinction also transferred across the expertise boundary: a model trained on the four traditions classified controls at 0.689, and a model trained on controls classified each tradition at 0.740–0.878 (Supplementary Fig. S3).

The cross-tradition test therefore provides the direct test of the shared-state question: a model trained without exposure to a given tradition can still distinguish meditation from task in that unseen tradition, and complexity is the leading contributor to this generalization.

### Robustness and exploratory analyses

The Med−Task distinction was reproduced by elastic-net logistic regression and Extra Trees classifiers refitted on the identical splits (Supplementary Fig. S5), indicating it is not specific to the Random Forest.

The feature families are not independent, which limits how far any effect can be attributed to a single family. Within each condition, the 21 measures carry only 3.4 effective dimensions of independent information. Summing the values of the separate categories gives 7.4, which overstates the total because it ignores the overlap between categories (Supplementary Fig. S4).

To test what this overlap means in practice, we let each category decode meditation versus task and tradition identity on its own, and compared its performance with that of random feature sets of the same size. Size alone accounts for a great deal: random sets of 2, 3, 7 and 9 features decoded meditation from task at 0.720, 0.764, 0.801 and 0.806 respectively, irrespective of which measures they contained. Against that reference, no family performed better than a random set of equal size.

Every category decoded both the state contrast (0.666–0.786) and tradition identity (0.343–0.574) above its own permutation chance level (*P*≤0.026), but none exceeded a random set of equal size on either axis (excess −0.101 to +0.026, all *P*>0.30), and only one of twelve pairwise comparisons survived FDR correction (*q*=0.024), close to what twelve tests would be expected to produce. Both the state change and the tradition signature are therefore encoded redundantly across the feature set, with contributions from multiple feature families (Supplementary Fig. S7).

Psychological ratings were collected only in the Isha Yoga study, so the exploratory brain–behaviour analysis is restricted to those participants. Within that cohort, the Med−Task change in Beta2 power correlated with self-rated concentration in controls (ρ=+0.55, n=17, *P*=0.023) but not in practitioners (ρ=−0.10, n=18, *P*=0.71). The two coefficients did not differ reliably (Fisher *z*=−1.86, *P*=0.063).

These relationships were selected from a grid of 7 bands × 6 scales, and none of the 84 correlations in that grid survives FDR correction. The full grid is shown in supplementary, with the reported correlations highlighted (Supplementary Figs. S6, S6b).

## Discussion

We set out to separate three things that are usually confounded in the meditation literature: what distinguishes meditation as a state, what changes with long-term practice, and what is specific to a tradition. Using a broad neurodynamic feature set across four traditions recorded against a common external task, three findings emerge.

Across four traditions, meditation was reliably separable from an external task, and the separation was driven mainly by a drop in the complexity of ongoing neural activity. All seven complexity measures decreased with large effects, and complexity carried the largest share of classifier. This meditation–task distinction was present in both practitioners and controls, and generalized to each tradition when that tradition was held out from training. Both long-term practice and technique then modify the shared profile. Practitioners and controls differed in the contribution of timescale features and in seven individual features. The four traditions differed more strongly still, in the contribution of the oscillatory, aperiodic and timescale families, in the Med−Task change of 18 of 21 features, and enough that an individual’s own profile carries recoverable information about which practice they follow.

Signal complexity was the only feature family whose contribution did not vary reliably across traditions, and it was the leading contributor in every leave-one-tradition-out model. A classifier that had never seen a tradition still separated meditation from task within it, in all four cases. This is a stronger claim than within-tradition accuracy can support, and it is the result on which the shared-state interpretation rests. Here, we use “shared state” to refer specifically to a neurodynamic pattern that distinguishes meditation from the external task and generalizes to a tradition that was not represented during training. Meditation, on this account, has a complexity-dominated core common to the practices we studied, overlaid by tradition-specific configurations of spectral, aperiodic and timescale activity. Given such importance of complexity dynamics, future studies could extend Topographic Reorganization model of Meditation (TRoM) to an integrated ‘Topographic-Dynamic Reorganization Model of Meditation’ ^12^.

Our finding of reduced complexity should be interpreted relative to external task state. Recent reviews report higher complexity during meditation and lower baseline complexity as a trait of practice, but the state comparisons underlying these findings have largely involved waking rest or mind-wandering ^19,20,22^. Rest and mind-wandering are internally generated states of relatively unconstrained content; whereas our comparator is an external task state. Our results characterize the axis of an internal–external dimension of neural dynamics ^9^, with the observed reduction representing lower complexity relative to external engagement and not necessarily an absolute reduction across all internally generated states. This suggests that complexity may increase along one axis (from rest to meditation) while decreasing along another (from meditation to external task). Future work should test this directly by considering all three conditions within the same participants.

What does such reduction of complexity at the core of the meditative state mean? It reflects more regular, lower-dimensional activity, in line with prior reports of reduced entropy in skilled practitioners ^21^. Both timescale measures increased during meditation, indicating longer autocorrelation and greater temporal continuity, alongside a steeper aperiodic slope, which has been linked to increased inhibitory tone ^23^.

The redundancy of the feature set constrains what “complexity-dominated” can mean. The seven complexity measures carry 1.21 effective dimensions and the battery as a whole about three, so any seven features span much the same space as complexity’s seven, and once feature count is held constant no family decodes either contrast better than an arbitrary set of the same size. Complexity’s prominence is therefore reliability and transferability, not exclusivity: it is the family the classifier weights most heavily, whose direction is invariant across traditions and expertise levels, and which dominates every leave-one-tradition-out model. This is a property of the measures rather than a limitation of our set: aperiodic slope, entropy, fractal dimension and high-frequency power are partly mathematical restatements of one another, since a flattening spectrum is also a more irregular signal. Category-level attributions common in this literature, that an effect is “a theta effect” or “a complexity effect”, are correspondingly weaker than usually presented. One asymmetry survives this levelling, with aperiodic features ranking first on the tradition axis and last on the state axis. This converges with the aperiodic slope carrying the strongest Group × Condition interaction across traditions.

Long-term practice left the main meditation signature intact: practitioners and controls showed the same decrease in complexity during meditation and differed reliably only in timescale features, which contributed less in practitioners. This is in line with recent evidence that intrinsic neural timescales may become less differentiated between internally and externally directed attention in advanced practitioners ^26^. We rest the remainder of the experience-level claim on the feature-level models rather than on category shares, since the complexity and oscillatory differences, though in the expected directions, did not survive correction. The aperiodic contribution was indistinguishable between groups, in line with a resting-state study finding differences in band-specific oscillatory power but not in aperiodic parameters ^32^. As a cross-sectional comparison, this design cannot establish that practice caused these differences.

The traditions differed markedly in which feature families carried their distinction, and we suggest, tentatively, that this reflects a correspondence between the attentional demands of each practice and the temporal structure of the accompanying neurodynamics. Isha Yoga is a breath-watching practice requiring sustained attention on the breath. Our findings show prominent contributions from oscillatory-rhythmic features in this tradition. We hypothesize that this may relate to the rhythmic nature of breath cycles, which has been shown to entrain cortical rhythms in other studies. Testing this hypothesis would require concurrent respiratory and EEG recording ^33–36^, consistent with the brain using intrinsic oscillatory dynamics to gate attention and sample information ^37^. Vipassana involves systematic body scanning, a temporally predictable attentional sequence; its similarly strong oscillatory contribution is compatible with the structured organization of sustained interoceptive attention, which likely engages insula and anterior cingulate regions supporting oscillatory activity related to body-state monitoring ^38^. The band involved matches the one previous multi-tradition comparison: Vipassana practitioners showed higher 7–11 Hz alpha activity than every other group examined ^15^, and alpha showed its largest Med−Task increase in Vipassana here. Both practices thus align with models of focused attention in which cyclic attention to bodily cues modulates ongoing neural oscillations ^3,39^, and that they are also the pair most often confused by the tradition classifier (Fig. 4b) is consistent with their sharing an attentional structure. This lends support to the assumption of shared temporal features at neural and mental levels, as recently postulated by the “theory of common currency” ^40,41^.

The other two practices depart from this pattern in complementary ways. Heartfulness involves sustained awareness directed toward the subjective heart space, facilitating receptive awareness and experiential absorption ^42,43^. As a continuous mode, it may favour temporal integration and stable neural dynamics, potentially contributing to the predominance of complexity and the uniquely large timescale contribution in its profile. This contrasts with Isha Yoga, in which timescale measures barely changed, consistent with evidence that intrinsic neural timescales can track differences across techniques ^27^. In Brahma Kumaris Raja Yoga, meditators visualize a soul centre between the eyebrows, a strategy centred on symbolic abstraction which makes it different from practices that focus on rhythmic bodily input. This is reflected in their profile which is dominated by aperiodic and complexity features. These practitioners are trained to meditate eyes-open ^44^ although recorded here eyes-closed, and that sensory–attentional dissociation should therefore be considered when interpreting its profile. Their training emphasizes rapid entry into meditative states, suggesting dynamics resembling non-linear transitions. These correspondences are interpretive and a future study could test with concurrent phenomenological and respiratory measurement.

The EEG signature of each tradition is reliable but modest compared with the signature shared by all four. Tradition identity was recoverable at 0.620 against a permutation chance of 0.245, well above chance but well below the 0.809 achieved for meditation versus task. That ordering is consistent with independent evidence that meditation styles are highly discriminable within an individual but generalize poorly across people, whereas the meditation-versus-non-meditation contrast is more robust ^45^. It also indicates that our leave-one-tradition-out result is not a trivial consequence of pooling: cross-population transfer of technique-specific information is precisely what tends to fail, and what transferred here was the state distinction rather than the technique.

### Limitations

The Brahma Kumaris Raja Yoga cohort behaved as an outlier throughout; its meditation epoch was also assembled from two one-minute segments of a protocol that alternated meditation with rest, whereas the task recording was continuous. However, such a splice-related concern could not have contributed to the autocorrelation and 1/f effects, as these were computed from 4-second sub epochs and were largest in this group. The tradition cohorts are small (n=25–48), so within-tradition accuracies show variability. The data comes from a single centre so multi-site replication, including non-Indian participants is needed to establish population-level generalizability. Thus, the concern that motivated the ENIGMA-Meditation consortium (Ganesan et al., 2025) applies here as to any single dataset. This is a strength as well, as all participants were recorded using the same acquisition system, task, and preprocessing pipeline, enabling harmonized cross-tradition comparisons from pan-India participants. We aggregated features across channels, so spatial heterogeneity in how meditation modulates different scalp regions is not captured. This precludes testing whether the shared complexity signature is spatially uniform or localized to specific networks (e.g., default-mode or task-positive regions). Future work with spatially resolved analysis is needed to integrate these findings with topographic models such as TRoM.

Taken together, these results support a shared neurodynamic signature of meditation relative to an external task, with diverse contemplative techniques showing both common and tradition-specific features. This shared signature is characterized primarily by reduced complexity, whereas tradition-specific profiles are shaped more strongly by oscillatory, aperiodic, and timescale features associated with the respective induction techniques. We acknowledge that our ’shared state’ claim is limited to the four traditions studied here, all of which are rooted in Indian contemplative traditions. Whether the complexity-dominated signature generalizes to other practices (e.g., Transcendental Meditation, Zen, loving-kindness) is unknown. Nevertheless, these complementary components provide a neurodynamic framework in which in which a common meditative signature coexists with tradition-specific signatures, with the latter sufficiently distinctive to identify an individual’s practice from their response profile. Our analyses use whole-brain averaged features, however, so the present findings do not establish a topographic organization of this shared or tradition-specific signature. Beyond meditation, the internal–external axis characterized here is a general dimension of brain organization, described most fully as the anticorrelated dynamics of default-mode and task-positive systems ^46^ and implicated across conditions from depressive rumination ^47^ to psychosis ^48^. That literature is almost entirely network- and fMRI-based; our findings show that this dimension also has a large and interpretable signature in the temporal statistics of EEG activity, one that transfers across contemplative techniques and levels of expertise. Future work combining these neurodynamic measures with spatially resolved EEG or other imaging modalities could test whether the shared and tradition-specific dynamics also have distinct topographic organization.

Future work combining these neurodynamic measures with spatially resolved EEG or other imaging modalities could test whether the shared and tradition-specific dynamics also have distinct topographic organization. Such work would provide a basis for extending the Topographic Reorganization Model of Meditation (TRoM) toward a topographic-dynamic framework ^12^.

## Methods

### Participants

We curated data from four EEG studies conducted at the same centre ^49–52^: practitioners of Brahma Kumaris Raja Yoga (BKY, n=48), Heartfulness (HFN, n=25), Isha Yoga (ISY, n=40) and Vipassana (VIP, n=43), and a pooled non-meditator control group (CNT, n=70), giving 226 participants.

Analyses of the paired meditation–task contrast use the 200 participants with usable data in both conditions. The participant sizes for each group in the paired analyses are available in Supplementary Table S3. Advanced meditators had extensive experience. Brahma Kumaris Raja Yoga practitioners had at least ten years of regular practice. Isha Yoga practitioners had a history equivalent to roughly 7–23 years of daily one-hour practice, including at least one advanced (“Samyama”) retreat.

Heartfulness practitioners had 6–28 years of practice. Vipassana participants were senior meditators or teachers with more than seven years of practice, including at least one advanced retreat. All participants were right-handed, in good general health, and screened for neurological or psychological disorders, substance abuse, and relevant medication. Meditators were excluded if they practiced any yoga or meditation outside their primary tradition (e.g., Vipassana practitioners who also practiced Transcendental Meditation). Practices within the same tradition (e.g., loving-kindness meditation for Vipassana practitioners) were not grounds for exclusion. Controls were excluded if they had any current or prior experience of either.

### Ethics

All participants gave written informed consent, and protocols were approved by the NIMHANS Institute Human Ethics Committee in accordance with the Declaration of Helsinki (1964).

### Design considerations and analytic constraints

We used an externally oriented task as the comparison condition as it provides a clearer contrast between externally and internally directed attention and a common reference point against which all four traditions can be compared. This choice is particularly relevant for experienced meditators, for whom resting-state activity may be altered by long-term practice and may contain substantial internally oriented or meditation-related activity, thereby reducing the interpretability of rest as a neutral comparison condition ^26,50,53^

The dataset combines four studies, so protocols differ as described below; all data were collected in the same laboratory with the same recording system and the same external task, and processed through a single pipeline. Group sizes are unequal, with Heartfulness the smallest (n=25).

Age differed across groups (Kruskal–Wallis *H*(4)=55.99, *P*<0.001) and was therefore treated as a group-level confound: mean-centred age was a fixed-effect covariate in every Group × Condition model, and its contribution to group identity was quantified by comparing EEG-only and EEG-plus-age versions of the composite classifier. Subject-level practice duration was not available across all studies, so experience is indexed by the categorical practitioner/control distinction.

The meditation condition was recorded with eyes closed and the task with eyes open. Alpha power is are sensitive to eye state independently of attentional orientation, so the state-level contrast is between an internally oriented eyes-closed state and an externally oriented eyes-open task. The contribution of alpha power to classification is reported as well. Tradition-level comparisons are unaffected, since all four traditions share the same contrast. Brahma Kumaris Raja Yoga practitioners were additionally recorded with eyes closed although their habitual practice is eyes-open (Sharma et al., 2023).

Features were aggregated across channels by trimmed mean, so spatial heterogeneity is not captured. Psychological rating scales were collected only in the Isha Yoga study; that analysis is restricted to those participants and reported as exploratory.

### Experimental protocol

For external attention, all studies used the same paradigm, ANGEL (Nair et al., 2016), a visual oddball task common to all studies. It provided a common externally oriented attentional comparison across all four meditation traditions. The task required attention to externally presented visual stimuli, providing a contrast with the internally oriented meditation conditions ^26^.

For internal attention, each meditation condition was rooted in its own tradition. Vipassana (40 minutes) involved focused attention with awareness of the impermanence of sensations (Kakumanu et al., 2018). Heartfulness (20 minutes) involved sustained attention to a sense of light within the heart (Sankar Sylapan et al., 2020). Isha Yoga (15 minutes) was a breath-watching practice (Malipeddi et al., 2024). Brahma Kumaris Raja Yoga (1 minute) involved soul-conscious meditation, in which participants attempted to experience the self as a point of light behind the forehead (Nair et al., 2017); controls otherwise focused on their breath.

### EEG acquisition and preprocessing

EEG was recorded with a Geodesic EEG System 300 (Electrical Geodesics Inc., USA), 128-channel HydroCel Sensor Nets and a Net Amps 300 amplifier, digitized at 24-bit resolution and 1 kHz, with impedance below 50 kΩ, at the Neurophysiology Department, NIMHANS, Bangalore. Stimuli were delivered via E-Prime 2 with time-locked event markers.

Preprocessing used MATLAB R2023b and EEGLAB v2023.1 ^54^ with a lab standardized pipeline applied uniformly across studies: band-pass (0.5–60 Hz) and notch (47–53 Hz) filtering, downsampling to 500 Hz, and bad-channel removal. This was followed by ICA-based artefact removal which used ICLabel with probability thresholds ranging from 0.99 to 0.60, with component removal capped at 10% of components and 30% of estimated brain signal. Artefact Subspace Reconstruction (15 SD), channel interpolation and common-average re-referencing were then applied. Two 30 second segments were visually inspected for residual artefacts, followed by further ICLabel-guided component removal where required. The central two minutes of each condition were analysed. This window length was chosen to balance signal stability (longer windows reduce variability in complexity and timescale estimates) against the constraint that the Brahma Kumaris Raja Yoga protocol provided only one-minute meditation segments. For that cohort, two one-minute segments were concatenated. As features were extracted from 4-second sub-epochs (see below), there could not have been any splice-related discontinuities. For the Brahma Kumaris Raja Yoga protocol, which had only one-minute meditation segments, two meditation segments were combined to match the duration with the other traditions. The central two-minute task segment was selected to match the meditation duration used for analysis across conditions. Feature extraction was performed on the full two-minute analysed segment after preprocessing.

### Feature extraction

Feature extraction used custom Python (3.10.0). Data were segmented into 4-second epochs with 2-second overlap. Features were computed for 51 scalp channels spanning frontal, frontocentral, central, centroparietal, parietal, parieto-occipital, and occipital regions (F7–F8, FC5–FC6, T7–T8, C5–C6, CP5–CP6, P7–P8, PO7–PO8, and O1–O2, including midline electrodes). These 51 channels were selected a priori to provide a common set of electrodes across datasets. Feature values were computed per epoch and channel and then aggregated across channels using a 10% trimmed mean. This aggregation discards spatial heterogeneity, so the present findings characterize whole-brain averaged dynamics for simpler interpretation of EEG dynamics and do not establish a topographic organization of the shared or tradition-specific signatures. Aperiodic and oscillatory features were derived via IRASA ^24^, using Welch power spectral density estimation over 1–50 Hz and resampling factors from 1.10 to 1.90 in steps of 0.05. The aperiodic component was estimated from the median across resampled spectra and fitted in log-power space to obtain the intercept, slope, and R². Oscillatory features included power in seven predefined bands: Delta (1–4 Hz), Theta (4–8 Hz), ThetaAlpha (6–10 Hz), Alpha (8–12 Hz), Beta1 (12–18 Hz), Beta2 (18–30 Hz), and Gamma1 (30–40 Hz) Additional oscillatory measures included the area under the oscillatory–aperiodic difference and the spectral edge of the oscillatory component. Complexity features and DFA used AntroPy ^55^. These comprised normalized permutation entropy, normalized SVD entropy, sample entropy, DFA, Petrosian, Katz, and Higuchi fractal dimensions, and normalized Lempel–Ziv complexity. Lempel–Ziv complexity was calculated after binarizing each signal at its within-epoch mean. All complexity and DFA measures were computed directly from the EEG time series. Autocorrelation window (ACW) was estimated from the autocorrelation function of each EEG epoch and channel as the first lag at which the autocorrelation fell to 0.50. In total, 21 features comprised four neurodynamic categories—oscillatory, aperiodic, complexity, and timescale (see Table 1).

### Classification

We use Med to denote the meditation condition and Task to denote the externally oriented ANGEL task. Med−Task denotes the within-subject difference between the meditation and task values for a given feature. Each classification observation corresponded to one subject-condition after epoch-level and channel-level aggregation. Random Forest classifiers ^62^ distinguished Med from Task using a subject-aware 80/20 train–test split stratified by source study. We selected Random Forest because it handles high-dimensional, correlated features without explicit regularization and provides interpretable feature importances via TreeSHAP. Robustness checks with elastic-net logistic regression and Extra Trees classifiers reproduced the main results (Supplementary Fig. S5). Hyperparameters were selected by 5-fold StratifiedGroupKFold grid search grouped by subject (*n_estimators* ∈ {100, 200}, *max_depth* ∈ {None, 10, 20}, *min_samples_leaf* ∈ {1, 3}), optimizing balanced accuracy. Median imputation and standardization were fitted on training data only. The procedure was repeated across 100 independent splits. Cohorts were: all participants (state level); practitioners and controls separately (experience level); and each tradition alone (tradition level). Controls are excluded from the tradition-level cohorts because that question concerns differences between techniques.

### Out-of-sample feature attribution

TreeSHAP ^63,64^ was applied to the held-out test subjects of each run, using the same fitted Random Forest that produced the reported accuracy, so attributions are cross-validated rather than fitted and explained on the same data. Each subject is held out in approximately 20 of 100 runs; their SHAP values were averaged to give one value per subject per condition, and the Med−Task contrast tested with a paired Wilcoxon signed-rank test across subjects with FDR correction.

SHAP values indicate how strongly a feature drives the classifier toward the meditation class, not whether the feature increases or decreases. For the binary meditation-versus-task classifier, a positive SHAP value indicates that a feature pushes the model’s prediction toward the meditation class, whereas a negative value pushes it toward the task class. SHAP magnitude therefore quantifies the feature’s contribution to the classification decision but does not by itself indicate the physiological direction of the feature change. Direction was therefore taken separately from the paired Cohen’s *d*z of the raw feature, and the two are presented as distinct quantities.

### Leave-one-tradition-out generalization

Within-tradition classification alone does not establish a shared neurodynamic signature, as separate classifiers could distinguish meditation from task using different features in each tradition. We therefore tested generalization across traditions by training classifiers on three traditions and testing them on the fully held-out fourth, repeated across 50 model seeds. Successful classification in the held-out tradition provides a key test of whether the meditation–task distinction generalizes across different contemplative techniques. Only advanced meditators were included in these analyses; controls were excluded from both the training and test sets. The null was constructed by permuting condition labels within subject in the training set (5,000 permutations). The observed value is averaged across seeds whereas each null value is a single fit, which makes the null slightly wider than the sampling distribution of the observed statistic and the test correspondingly conservative. Confidence intervals were obtained by bootstrap resampling of held-out subjects from the existing predictions, which reflects participant-level uncertainty; the spread across model seeds reflects only stochastic variation and is not used as an error bar.

### Category-level importance and subject-level inference

Feature importance was quantified using TreeSHAP values from the held-out Random Forest predictions. Feature importances were summed within each category and expressed as shares. As the 100 runs are repeated resamplings of the same participants rather than independent observations, these distributions are reported descriptively. Inference uses the subject as the resampling unit, with bootstrap confidence intervals over subjects (2,000 replicates) and permutation tests shuffling group or tradition labels across subjects (5,000 permutations). The statistic was the between-group difference or, for the four traditions, the spread across traditions.

As the categories differ in size and internal redundancy, comparisons were repeated with importance expressed per feature and per effective dimension. Effective dimension was estimated from the participation ratio of the within-category Spearman correlation matrix computed once on the full paired cohort and reused throughout. The correlation matrix was not re-estimated within subgroups because the smaller sample sizes would make these estimates less stable. The effective-dimension adjustment changes the statistic and is not simply a rescaling of the raw importance share.

Permutation tests were therefore repeated using the adjusted shares. The same participation ratio applied to all 21 features gives the global effective dimensionality. It was calculated both pooled across conditions and separately within each condition. The two approaches gave similar results, indicating that the observed redundancy was not driven by the state contrast.

### Mixed-effects models

For each of the 21 features, rank-normalized via the van der Waerden transformation, we fitted: feature ∼ Group × Condition + centred Age + (1 | Subject). The interaction was tested by likelihood-ratio test against the model without it, with age retained in both, and FDR correction across features. Group comprised practitioners and controls at the experience level, and the four traditions at the tradition level. Estimated Med−Task changes are model-derived marginal predictions at mean age.

### Composite classifier

Subject-level Med−Task deltas across the 21 features were used to decode group identity with a consensus of Random Forest and Extra Trees classifiers, evaluated by repeated stratified 5-fold cross-validation, with bootstrap confidence intervals. The permutation null was generated through the identical procedure, so the comparison against chance is unaffected. Due to unequal group sizes, chance level was derived by permuting group labels across subjects. Four-class (traditions) and five-class (traditions plus controls) versions were run with and without age.

### Category-restricted decoding

To test whether any feature family carries information the others do not, each category decoded on its own on two axes: meditation versus task using one observation per subject per condition after epoch- and channel-level aggregation, and tradition identity from subject-level Med−Task deltas. Cross-validation folds were grouped by subject throughout; during bootstrap resampling, grouping used the original subject identifier so that duplicated participants could not appear in both training and test folds. Each category used all of its features and was referenced to a null of 200 random feature sets of the same size drawn from all 21 features, giving a size-controlled excess comparable across categories of different sizes. Chance levels came from 1,000 permutations of the class label, across subjects for the tradition axis and within subject for the state axis. Confidence intervals are from 500 bootstrap resamples of subjects, and comparisons between categories were computed on the same resamples, so the contrast is paired, and on the size-controlled excess rather than on raw accuracy. The random sets were drawn from the whole battery, including the category’s own features, making the reference conservative. A random seven-feature set contains 2.3 complexity measures on average. This analysis was is reported as exploratory.

### Robustness

Elastic-net logistic regression and Extra Trees classifiers were refitted on the identical train/test splits used by the Random Forest. These analyses assessed whether the meditation–task distinction was reproducible with different classifier families.

### Exploratory brain–behaviour analysis

For Isha Yoga participants with post-session ratings, subject-level Med−Task deltas were correlated with rating scores using Spearman correlations with bootstrap confidence intervals, computed separately in practitioners and controls, and compared with a Fisher r-to-z test using the variance appropriate to Spearman coefficients. The reported relationships were identified within a grid of 7 bands × 6 scales computed in both groups (84 correlations). Whole-grid FDR-corrected values are therefore reported alongside uncorrected values.

## Data and Code Availability

The datasets and code are available from the corresponding author on reasonable request, subject to the data-sharing conditions of the ethics approval under which they were collected.

## Acknowledgements

We express gratitude to Ratna Jyothi and Bhuvanesh S. Sylapan who contributed to the original datasets used in this study. This work was supported by the National Institute of Mental Health and Neuro Sciences (NIMHANS) through institutional fellowships and research facilities. We gratefully acknowledge funding from the Cognitive Science Research Initiative (DST-CSI), Department of Science & Technology, Government of India, New Delhi (Grant No. SR/CSI/63/2011 to B.M.K.), and a six-month fellowship provided by the Heartfulness Institute. None of the funding agencies had any involvement in the research design, data collection, analysis, interpretation, or the decision to publish this work.

## Supplementary Information

Supplementary file is available in a PDF format.

## Contributions

Conceptualization: P.S., A.S., S.M., R.P.N., B.M.K. and G.N.; Data curation: P.S. and A.S.; Formal analysis: P.S, and A.S.; Methodology: P.S., A.S., S.M., R.V.G.; Additional Discussions: B.V., A.K.N, P.K.BT.; Resources: B.M.K. and G.N.; Software, P.S. and A.S.; Supervision: A.S., R.P.N, G.N. and B.M.K.; Validation: A.S., S.M and R.V.G; Visualization: P.S. and B.V.; Writing – original draft: P.S., A.S., S.M., G.N.; Writing – review & editing: R.V.G, R.P.N., B.V., A.K.N., P.K.B.T., B.M.K, and G.N.

