## Supplementary file for "Distinct contemplative traditions share a common EEG complexity signature"

#### Author Information

Prakash Shrimali[1,2*], Arun Sasidharan[1*#], Saketh Malipeddi[2,] Bianca Ventura[2], Rahul Venugopal[1], Ajay Kumar Nair[3], Prejaas K.B. Tewarie[4], Ravindra P.N.[1], Bindu M Kutty[1#], Georg Northoff[2*#]

Affiliations

**[1]Centre for Consciousness Studies (CCS), Department of Neurophysiology, Basic Science Block, National Institute of Mental Health and Neuro Sciences (NIMHANS), Bengaluru, 560029, Karnataka, India**

**[2]The Royal's Institute of Mental Health Research & University of Ottawa, Brain and Mind Research Institute, Centre for Neural Dynamics, Faculty of Medicine, University of Ottawa, 145 Carling Avenue, Rm. 6435, Ottawa, ON K1Z 7K4, Canada; Department of Cellular and Molecular Medicine, University of Ottawa, Ottawa, ON, Canada.**

**[3]Centre for Brain and Mind (CBM), Department of Psychiatry, National Institute of Mental Health and Neuro Sciences (NIMHANS), Bengaluru, Karnataka, India.**

**[4]Joint International Research Unit on Consciousness, CERVO Brain Research Centre, University of Laval, Québec City, G1J 2G3, Canada; Sir Peter Mansfield Imaging Centre, School of Physics, University of Nottingham, Nottingham, NG7 2RD, United Kingdom**

*Corresponding authors

[#]These authors jointly supervised this work

#### Supplementary Information

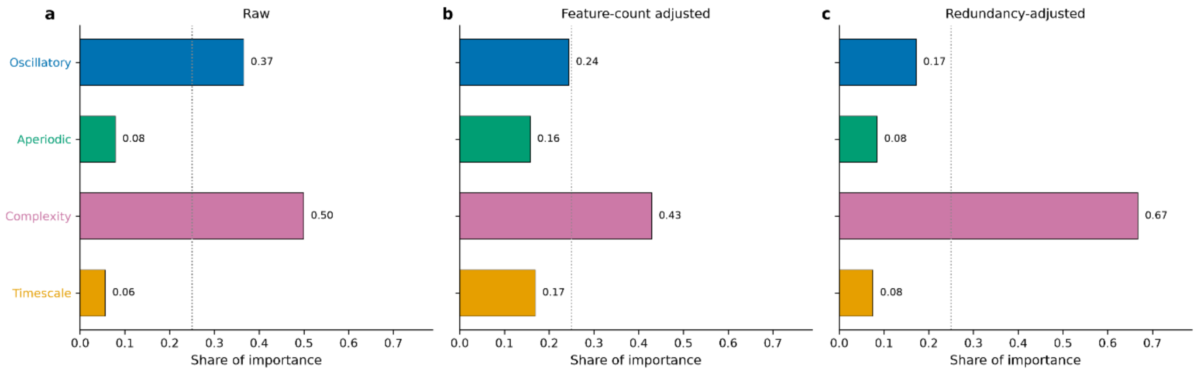


**Supplementary Fig. S1 | Category dominance under three normalizations.** Share of classifier importance per feature family in the pooled cohort, expressed as raw shares, per feature, and per effective dimension. Complexity leads under all three, so its contribution is not explained by the number or redundancy of its measures.

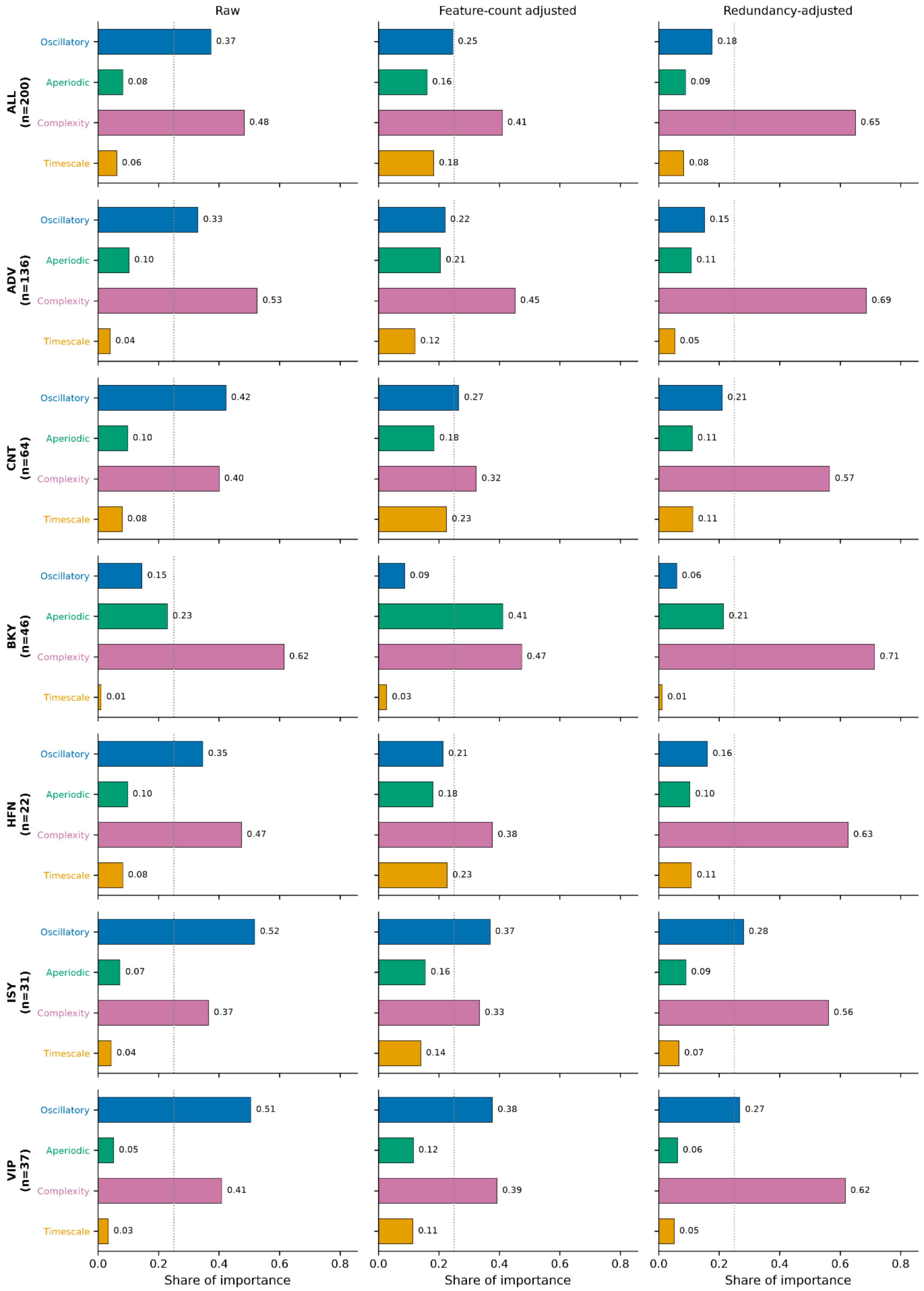


**Supplementary Fig. S1b | The same three normalizations within each group and tradition.** Rows are the pooled cohort, the two experience-level groups and the four traditions; columns are the three weightings. The dotted line marks an equal share across four families. In the control, Isha Yoga and Vipassana cohorts the leading family changes with weighting, raw shares placing oscillatory features first and both adjusted weightings placing complexity first.

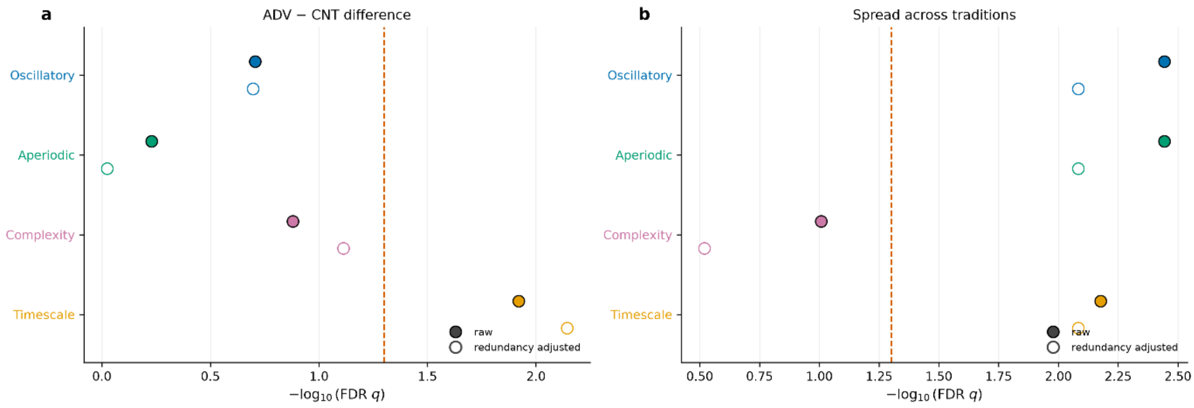


**Supplementary Fig. S1c | Category-level permutation results under raw and redundancy-adjusted weighting.** FDR-adjusted significance of the category-level permutation tests computed on raw shares (filled) and on shares per effective dimension (open). Because the adjustment renormalizes across families it is not a monotone rescaling of the raw share, so each test was re-run rather than inherited. Dashed line marks $q$=0.05. **a,** Difference between practitioners and controls. **b,** Spread across the four traditions.

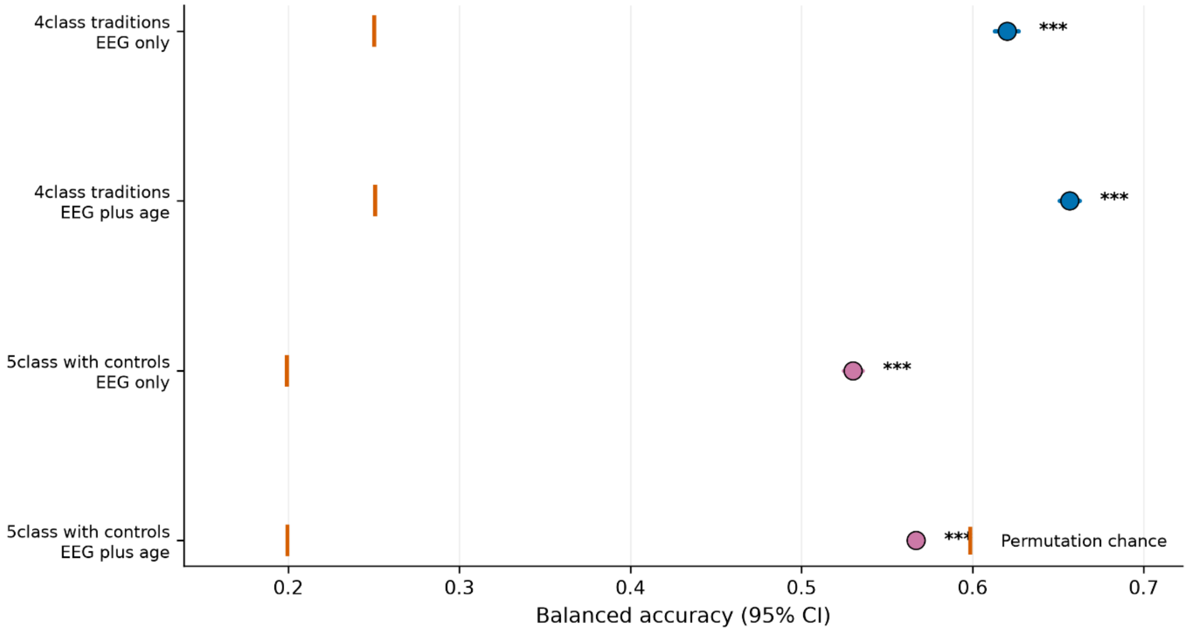


**Supplementary Fig. S2 | Composite decoding across configurations.** Balanced accuracy for decoding group identity from each participant's Med−Task delta profile, for the four-class (traditions only) and five-class (traditions plus controls) problems, with and without age as a predictor, against permutation-derived chance levels.

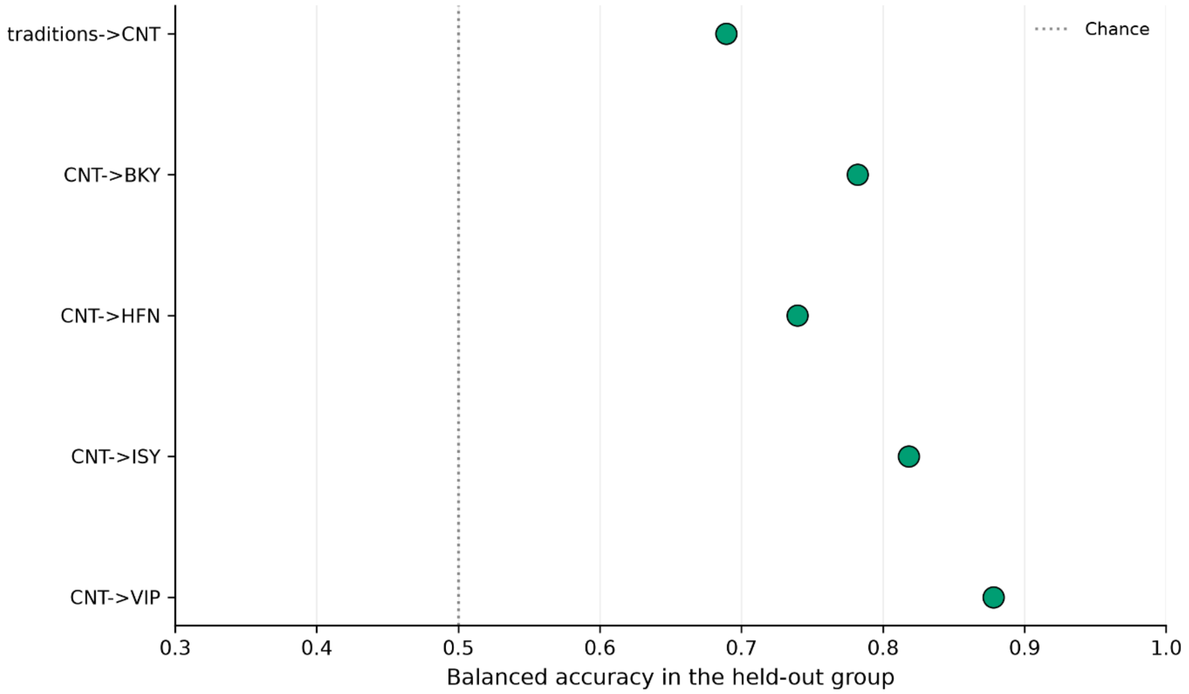


**Supplementary Fig. S3 | Cross-expertise transfer.** Balanced accuracy when a model trained on the four traditions is tested on controls, and when a model trained on controls is tested on each tradition

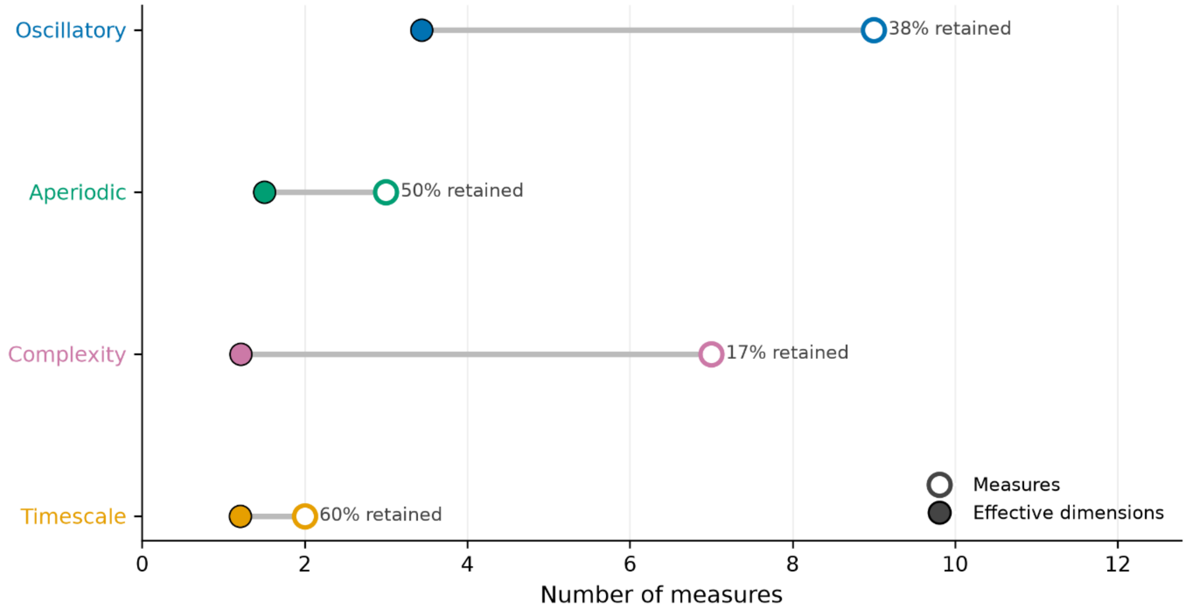


**Supplementary Fig. S4 | Within-category and global redundancy.** Effective dimensionality of each feature family from the participation ratio of its within-family Spearman correlation matrix, with

the mean absolute correlation among its measures, and the equivalent statistic computed across all 21 features pooled across conditions and within each condition separately.

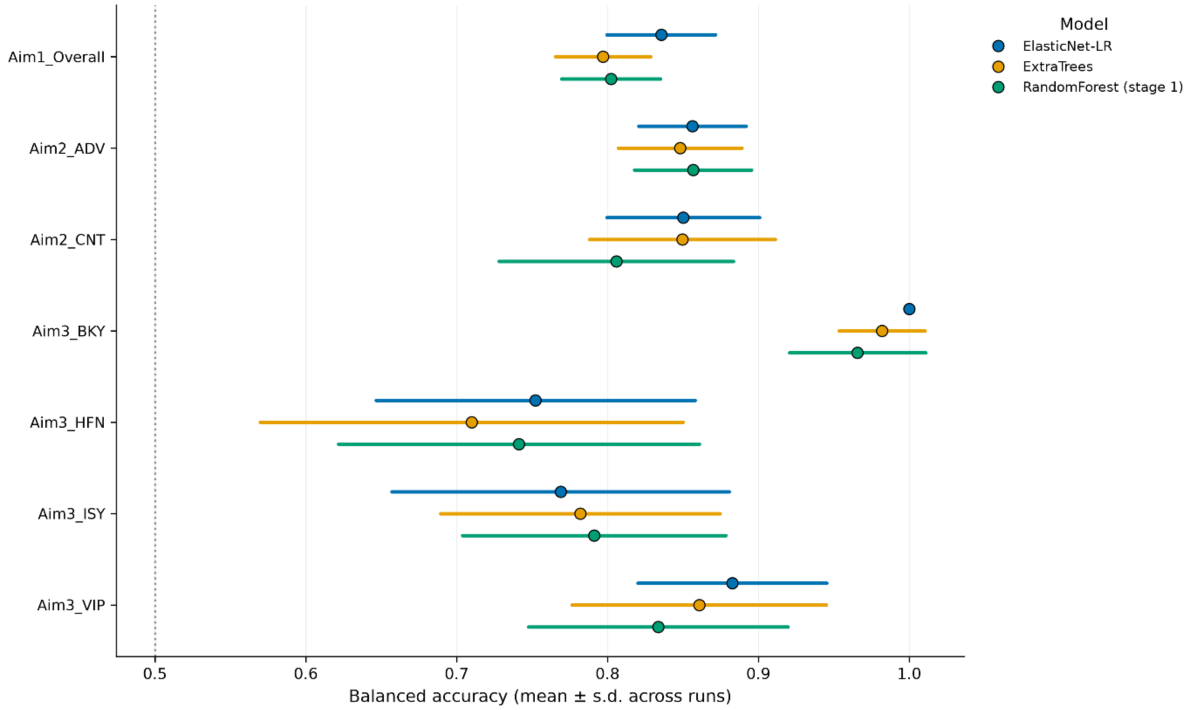


**Supplementary Fig. S5 | Model-family robustness.** Balanced accuracy of elastic-net logistic regression and Extra Trees refitted on the same train/test splits used in the main classification, alongside the Random Forest result, for each cohort.

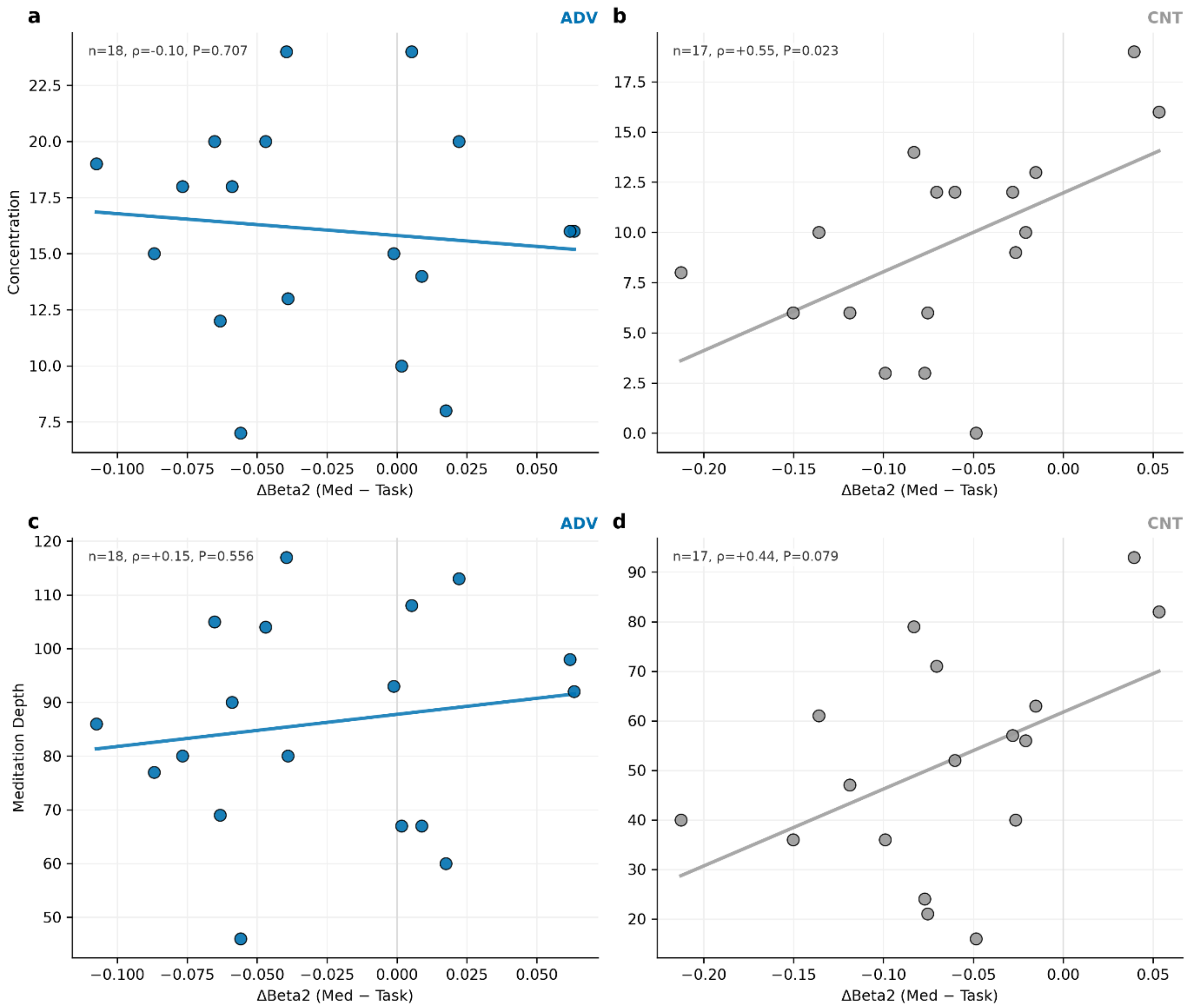


**Supplementary Fig. S6 | Exploratory Beta2 by rating scatter.** Subject-level Med−Task change in Beta2 power against self-rated concentration and meditation depth, separately for practitioners and controls, in the Isha Yoga cohort where ratings were collected. Lines are ordinary least squares fits shown for orientation only; statistics are Spearman correlations.

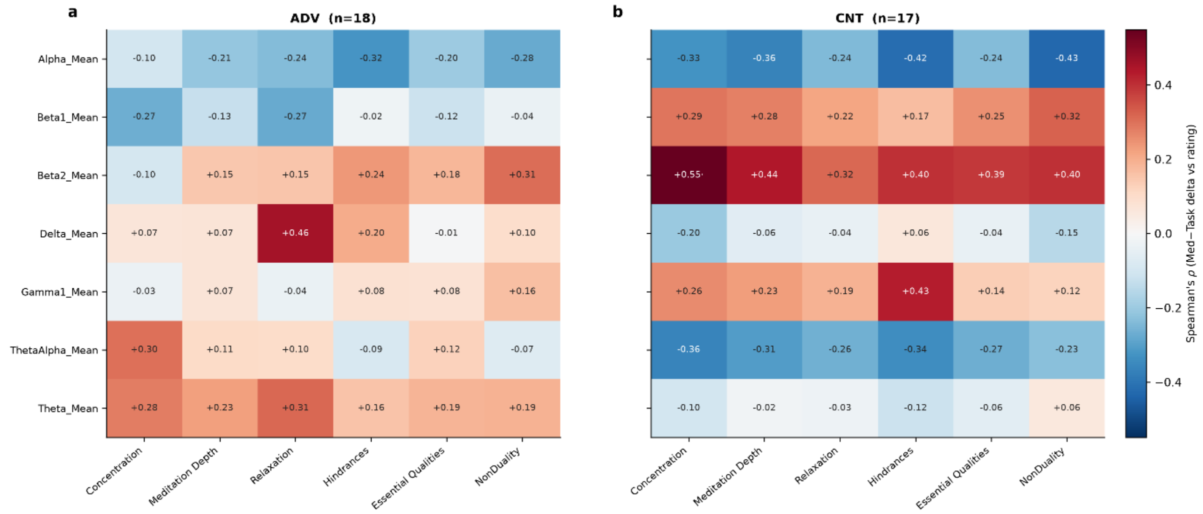


**Supplementary Fig. S6b | The full exploratory correlation grid.** Spearman correlations between each oscillatory band's Med−Task change and each post-session rating, shown separately for practitioners and controls. All 84 correlations are displayed. Asterisks mark cells surviving FDR correction across the entire grid; interpuncts mark cells reaching *P*<0.05 uncorrected only. Outlined cells are the focal comparisons shown in Supplementary Fig. S6. The grid is reported in full so that the focal panels are read against the multiplicity they sit in.

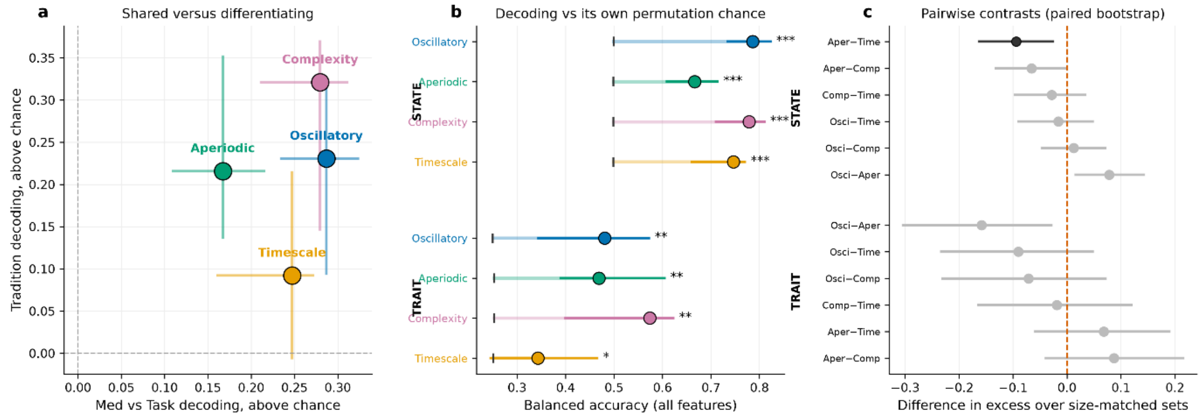


**Supplementary Fig. S7 | No feature family carries information the others do not.** Each family decodes on its own using all of its features, referenced to a null of random feature sets of the same size drawn from the full battery, so families of different sizes are comparable. **a,** Families positioned by how well they decode the meditation–task state change (x) and tradition identity (y), each relative to its own permutation chance level; bars are 95% subject-bootstrap intervals. Positions on these axes are not size-controlled and are descriptive only. **b,** Balanced accuracy against the permutation chance level for that family (tick), with 95% subject-bootstrap intervals; stars mark FDR $q<0.05$ against the permutation null. Seven of the eight $P$ values are at the permutation floor, so differences in the number of asterisks between the two axes reflect FDR correction within axis rather than differences in evidence. **c,** Differences between families in excess over size-matched random sets, computed on the same bootstrap resamples so the comparison is paired, FDR-corrected within axis; dark markers are $q<0.05$. Intervals are uncorrected, so an interval excluding zero with a grey marker denotes a nominally significant difference that does not survive correction. Only one of twelve contrasts reached significance, so the ordering is descriptive: all four families carried both shared and tradition-specific information, with none statistically dominating either axis.

### Supplementary Tables

**Supplementary Table S1 | Within-family and global effective dimensionality.**

| Feature family | Measures ($k$) | Effective dimensions | % independent | Mean abs. correlation |
|---|---|---|---|---|
| Oscillatory | 9 | 3.44 | 38 | 0.39 |
| Aperiodic | 3 | 1.50 | 50 | 0.70 |
| Complexity | 7 | 1.21 | 17 | 0.89 |
| Timescale | 2 | 1.21 | 60 | 0.81 |
| All features, conditions pooled | 21 | 2.83 | 13 | 0.50 |
| All features, meditation only | 21 | 3.42 | 16 | 0.44 |
| All features, task only | 21 | 2.93 | 14 | 0.48 |

Summing the within-family values gives 7.36, an upper bound that ignores redundancy between families. Pooled and within-condition estimates agree, indicating the redundancy is a property of the measures rather than of the state contrast.

**Supplementary Table S2 | Category importance under three normalizations, pooled cohort.**

| Feature family | Raw share | Per feature | Per effective dimension |
|---|---|---|---|
| Oscillatory | 0.365 | 0.244 | 0.172 |
| Aperiodic | 0.079 | 0.158 | 0.085 |
| Complexity | 0.500 | 0.429 | 0.668 |
| Timescale | 0.056 | 0.169 | 0.075 |

**Supplementary Table S3 | Bootstrap category shares by cohort, raw (2,000 replicates).** Values are shares of classifier importance with 95% confidence intervals.

| Cohort | *n* | Oscillatory | Aperiodic | Complexity | Timescale |
|---|---|---|---|---|---|
| All participants | 200 | 0.373 [0.313, 0.429] | 0.082 [0.070, 0.103] | 0.483 [0.423, 0.553] | 0.062 [0.046, 0.072] |
| Practitioners | 136 | 0.330 [0.259, 0.395] | 0.103 [0.067, 0.129] | 0.527 [0.460, 0.614] | 0.040 [0.030, 0.059] |
| Controls | 64 | 0.423 [0.332, 0.546] | 0.098 [0.070, 0.140] | 0.400 [0.280, 0.517] | 0.080 [0.045, 0.090] |
| Brahma Kumaris Raja Yoga | 46 | 0.146 [0.067, 0.219] | 0.229 [0.161, 0.313] | 0.615 [0.504, 0.714] | 0.010 [0.005, 0.035] |
| Heartfulness | 22 | 0.346 [0.207, 0.506] | 0.097 [0.063, 0.188] | 0.474 [0.273, 0.631] | 0.082 [0.044, 0.150] |
| Isha Yoga | 31 | 0.518 [0.327, 0.654] | 0.073 [0.045, 0.114] | 0.365 [0.245, 0.541] | 0.044 [0.024, 0.076] |
| Vipassana | 37 | 0.505 [0.307, 0.638] | 0.052 [0.026, 0.095] | 0.409 [0.272, 0.607] | 0.034 [0.013, 0.058] |

**Supplementary Table S3b | The same shares expressed per effective dimension.** Redundancy-adjusted shares with 95% confidence intervals.

| Cohort | *n* | Oscillatory | Aperiodic | Complexity | Timescale |
|---|---|---|---|---|---|
| All participants | 200 | 0.177 [0.141, 0.214] | 0.089 [0.075, 0.112] | 0.651 [0.596, 0.706] | 0.084 [0.061, 0.098] |
| Practitioners | 136 | 0.152 [0.112, 0.191] | 0.108 [0.070, 0.135] | 0.687 [0.631, 0.756] | 0.053 [0.038, 0.077] |
| Controls | 64 | 0.210 [0.152, 0.306] | 0.111 [0.079, 0.163] | 0.565 [0.443, 0.674] | 0.113 [0.063, 0.131] |
| Brahma Kumaris Raja Yoga | 46 | 0.060 [0.026, 0.095] | 0.215 [0.151, 0.294] | 0.714 [0.614, 0.789] | 0.012 [0.005, 0.040] |
| Heartfulness | 22 | 0.161 [0.087, 0.273] | 0.104 [0.065, 0.214] | 0.627 [0.415, 0.750] | 0.109 [0.058, 0.196] |
| Isha Yoga | 31 | 0.281 [0.149, 0.408] | 0.090 [0.055, 0.135] | 0.562 [0.432, 0.710] | 0.068 [0.038, 0.112] |
| Vipassana | 37 | 0.269 [0.136, 0.390] | 0.063 [0.031, 0.112] | 0.617 [0.464, 0.779] | 0.052 [0.019, 0.090] |

Complexity leads in every cohort once expressed per effective dimension, including the control, Isha Yoga and Vipassana cohorts, in which raw shares place oscillatory features first.

**Supplementary Table S4 | Leave-one-tradition-out generalization.** A model trained on three traditions tested on the fully held-out fourth. Permutation nulls were 0.500–0.501 in all four cases.

Seed SD, reflecting model stochasticity on a fixed test set, was 0.007–0.012; the bootstrap interval is the participant-level uncertainty and is the one plotted.

| Held-out tradition | *n* | Bal. accuracy | 95% CI | AUC | *P* | Leading family |
|---|---|---|---|---|---|---|
| Brahma Kumaris | 46 | 0.886 | [0.826, 0.946] | 0.935 | 0.0002 | Complexity (0.443) |
| Heartfulness | 22 | 0.707 | [0.591, 0.818] | 0.779 | 0.0086 | Complexity (0.526) |
| Isha Yoga | 31 | 0.821 | [0.742, 0.903] | 0.879 | 0.0002 | Complexity (0.557) |
| Vipassana | 37 | 0.852 | [0.783, 0.919] | 0.901 | 0.0002 | Complexity (0.501) |

**Supplementary Table S5 | Cross-expertise transfer.**

| Train | Test | Balanced accuracy |
|---|---|---|
| Four traditions | Controls | 0.689 |
| Controls | Brahma Kumaris Raja Yoga | 0.782 |
| Controls | Heartfulness | 0.740 |
| Controls | Isha Yoga | 0.818 |
| Controls | Vipassana | 0.878 |

**Supplementary Table S6 | Age-adjusted Group × Condition interactions.** Likelihood-ratio tests against a model without the interaction, on rank-normalized features with a random intercept per participant and mean-centred age in both models. FDR-corrected across the 21 features within each analysis.

| Analysis | Feature | LR | *q* |
|---|---|---|---|
| Practitioners vs controls (7/21 significant) | Permutation entropy | 12.61 | $4.13 \times 10^{-3}$ |
| | Petrosian fractal dimension | 12.57 | $4.13 \times 10^{-3}$ |
| | Gamma1 power | 8.76 | $1.81 \times 10^{-2}$ |
| | Aperiodic slope | 8.56 | $1.81 \times 10^{-2}$ |
| | Oscillatory area | 7.45 | $2.67 \times 10^{-2}$ |
| | Higuchi fractal dimension | 6.76 | $3.27 \times 10^{-2}$ |
| | Delta power | 6.41 | $3.40 \times 10^{-2}$ |
| Four traditions (18/21 significant; eight largest shown) | Aperiodic slope | 123.27 | $3.20 \times 10^{-25}$ |
| | Petrosian fractal dimension | 63.58 | $7.60 \times 10^{-13}$ |
| | Permutation entropy | 63.43 | $7.60 \times 10^{-13}$ |

| | | | |
|---|---|---|---|
| | Gamma1 power | 48.10 | 1.06 × $10^{-9}$ |
| | Aperiodic intercept | 32.55 | 1.69 × $10^{-6}$ |
| | Higuchi fractal dimension | 27.01 | 2.05 × $10^{-5}$ |
| | Aperiodic fit ($R^2$) | 26.45 | 2.30 × $10^{-5}$ |
| | Theta/Alpha ratio | 17.88 | 1.06 × $10^{-3}$ |

**Supplementary Table S7 | Composite decoding of group identity from Med−Task profiles.**

| Problem | Predictors | Balanced accuracy | 95% CI | Naive chance | Permutation chance | *P* |
|---|---|---|---|---|---|---|
| Four traditions | EEG only | 0.620 | [0.613, 0.627] | 0.250 | 0.250 | <0.001 |
| Four traditions | EEG + age | 0.657 | [0.651, 0.663] | 0.250 | 0.251 | <0.001 |
| Five classes with controls | EEG only | 0.530 | [0.525, 0.536] | 0.200 | 0.199 | <0.001 |
| Five classes with controls | EEG + age | 0.567 | [0.562, 0.571] | 0.200 | 0.200 | <0.001 |

Four-class confusion matrix, row-normalized and out-of-fold:

| True Predicted | BKY | HFN | ISY | VIP |
|---|---|---|---|---|
| Brahma Kumaris Raja Yoga | 0.96 | 0.00 | 0.02 | 0.03 |
| Heartfulness | 0.05 | 0.48 | 0.13 | 0.34 |
| Isha Yoga | 0.09 | 0.09 | 0.45 | 0.37 |
| Vipassana | 0.01 | 0.15 | 0.25 | 0.59 |

**Supplementary Table S8 | Model-family robustness.** Balanced accuracy, mean and SD across the 25 splits retained from the main classification, refitted on identical train/test partitions.

| Cohort | Elastic-net | Extra Trees | Random Forest |
|---|---|---|---|
| All participants | 0.836 (0.036) | 0.797 (0.032) | 0.802 (0.033) |
| Practitioners | 0.856 (0.036) | 0.848 (0.041) | 0.857 (0.039) |
| Controls | 0.850 (0.051) | 0.850 (0.062) | 0.806 (0.078) |
| Brahma Kumaris Raja Yoga | 1.000 (0.000) | 0.982 (0.028) | 0.966 (0.045) |
| Heartfulness | 0.752 (0.106) | 0.710 (0.140) | 0.741 (0.120) |
| Isha Yoga | 0.769 (0.112) | 0.782 (0.093) | 0.791 (0.087) |
| Vipassana | 0.883 (0.063) | 0.861 (0.084) | 0.834 (0.086) |

**Supplementary Table S9 | Category-restricted decoding.** Each family decodes alone using all of its features (*k* given). The size-matched reference is the mean accuracy of 200 random feature sets of the same size drawn from all 21 features. *P*(label) tests against the permutation null; *P*(size) against the size-matched null. Permutation chance was 0.250–0.253 on the tradition axis and 0.499 on the state axis.

*Tradition axis (which of four traditions; 136 practitioners).*

| Family | *k* | Bal. accuracy | 95% CI | Size-matched ref. | Excess | *P* (label) | *P* (size) |
|---|---|---|---|---|---|---|---|
| Oscillatory | 9 | 0.481 | [0.344, 0.572] | 0.582 | −0.101 | 0.001 | 0.990 |
| Aperiodic | 3 | 0.469 | [0.390, 0.604] | 0.452 | +0.017 | 0.001 | 0.413 |
| Complexity | 7 | 0.574 | [0.400, 0.622] | 0.560 | +0.014 | 0.001 | 0.433 |
| Timescale | 2 | 0.343 | [0.245, 0.465] | 0.389 | −0.045 | 0.026 | 0.726 |

*State axis (meditation versus task; 200 participants).*

| Family | *k* | Bal. accuracy | 95% CI | Size-matched ref. | Excess | *P* (label) | *P* (size) |
|---|---|---|---|---|---|---|---|
| Oscillatory | 9 | 0.786 | [0.734, 0.823] | 0.806 | −0.020 | 0.001 | 0.930 |
| Aperiodic | 3 | 0.666 | [0.609, 0.714] | 0.764 | −0.097 | 0.001 | 0.995 |
| Complexity | 7 | 0.779 | [0.711, 0.810] | 0.801 | −0.022 | 0.001 | 0.955 |
| Timescale | 2 | 0.746 | [0.660, 0.770] | 0.720 | +0.026 | 0.001 | 0.304 |

No family exceeded a random set of equal size on either axis.

**Supplementary Table S10 | Pairwise family contrasts on size-controlled excess.** Differences computed on shared bootstrap resamples, so the comparison is paired. FDR-corrected within axis.

| Axis | Contrast | Difference | 95% CI | *q* |
|---|---|---|---|---|
| State | Timescale − Aperiodic | +0.094 | [+0.027, +0.163] | **0.024** |
| State | Oscillatory − Aperiodic | +0.078 | [+0.017, +0.142] | 0.072 |
| State | Complexity − Aperiodic | +0.066 | [+0.002, +0.131] | 0.096 |
| State | Complexity − Timescale | −0.028 | [−0.096, +0.034] | 0.528 |
| State | Oscillatory − Timescale | −0.016 | [−0.089, +0.047] | 0.680 |
| State | Oscillatory − Complexity | +0.012 | [−0.046, +0.071] | 0.680 |
| Tradition | Aperiodic − Oscillatory | +0.158 | [+0.029, +0.303] | 0.096 |
| Tradition | Complexity − Oscillatory | +0.071 | [−0.072, +0.230] | 0.480 |

| Axis | Contrast | Difference | 95% CI | *q* |
|---|---|---|---|---|
| Tradition | Aperiodic − Complexity | +0.087 | [−0.039, +0.215] | 0.424 |
| Tradition | Timescale − Oscillatory | +0.090 | [−0.048, +0.233] | 0.424 |
| Tradition | Aperiodic − Timescale | +0.068 | [−0.058, +0.189] | 0.462 |
| Tradition | Complexity − Timescale | +0.019 | [−0.119, +0.164] | 0.784 |

Intervals are uncorrected; only the first row survives FDR correction.

**Supplementary Table S11 | Exploratory brain–behaviour correlations.** Isha Yoga cohort, the only study in which post-session ratings were collected. Med−Task change in Beta2 power against each rating.

| Rating | Group | *n* | Spearman ρ | *P* | 95% CI |
|---|---|---|---|---|---|
| Concentration | Practitioners | 18 | −0.095 | 0.707 | [−0.500, +0.333] |
| Concentration | Controls | 17 | +0.549 | 0.023 | [+0.076, +0.839] |
| Meditation depth | Practitioners | 18 | +0.149 | 0.556 | [−0.336, +0.551] |
| Meditation depth | Controls | 17 | +0.438 | 0.079 | [−0.080, +0.780] |

Between-group comparison of these correlations: concentration, Fisher $z=-1.861$, $P=0.063$; meditation depth, Fisher $z=-0.836$, $P=0.403$. The two groups do not differ reliably in either. These cells were identified within the full 84-correlation grid (Supplementary Fig. S6b), in which no correlation survives FDR correction across the grid.
